# A stress-responsive morphogenetic program coordinates uterine luminal remodeling during early pregnancy

**DOI:** 10.64898/2026.01.29.702684

**Authors:** Chihiro Ishizawa, Shizu Aikawa, Yamato Fukui, Xueting He, Ryoko Shimizu-Hirota, Daiki Hiratsuka, Mitsunori Matsuo, Takehiro Hiraoka, Yasushi Hirota

## Abstract

Successful embryo implantation requires coordinated interactions among the endometrial epithelium, stroma, and the embryo, yet the underlying mechanisms have not been fully understood. Using three-dimensional histological reconstruction combined with single-cell and spatial transcriptomics, we identify a previously unrecognized phase of luminal architectural reorganization preceding embryo attachment. Within a narrow peri-implantation window, the luminal epithelium rapidly remodels from a highly folded structure into a flattened, organized architecture that is spatially associated with embryo positioning. This morphogenetic transition is accompanied by activation of stress-responsive signaling across the epithelial and stromal compartments. Functional analyses show that uterine-specific deletion of the stress-responsive MAP kinase p38α disrupts luminal remodeling, leading to persistent epithelial folding, failed embryo attachment, and infertility despite normal hormone levels and embryo development. In contrast, epithelial-specific deletion of p38α causes only mild uterine abnormalities, indicating that epithelial p38α loss alone is insufficient to account for the severe implantation phenotype. Although combined progesterone and leukemia inhibitory factor supplementation rescues embryo attachment in p38α-deficient uteri, luminal disorganization, abnormal stromal responses, and impaired pregnancy progression persist. These findings identify a uterine p38α-dependent, stress-responsive program that supports luminal remodeling and epithelial-stromal communication required for implantation competence.

## INTRODUCTION

Infertility is a social concern that affects 17.5% of the adult population worldwide^1^. Some patients who undergo *in vitro* fertilization and embryo transfer (IVF-ET) repeatedly fail to become pregnant, even after ET using high-quality embryos^2^. Embryo implantation, which is the starting point of pregnancy, can be divided into several processes, including blastocyst spacing, apposition, attachment to the uterine luminal epithelium, and invasion into the endometrial stroma^2, 3^. Establishing implantation requires an exquisite interaction between the endometrium and the embryo; however, the detailed underlying mechanisms remain unclear.

Based on the similarities in the influence of female sex hormones on the endometrium during pregnancy, rodents have been utilized as an *in vivo* model of human pregnancy. In mice, day 1 of pregnancy is defined by the observation of a vaginal plug. After coitus, high serum levels of estrogen (E₂) induce epithelial proliferation, and P_4_ production from the ovary increases on day 3. Embryos in the oviduct reach the uterus on day 4 in a P_4_-dominant hormonal environment. In the uterus, proliferation–differentiation switching (PDS), characterized by the suppression of epithelial proliferation and the induction of stromal proliferation, occurs on day 4 under the continuous influence of P_4_. This transition enables the endometrium to acquire receptivity for embryo implantation^4, 5^. Along with this dynamic change, the morphology of the endometrial luminal epithelium exhibits a slit-like narrowing, known as the formation of a slit-like luminal structure^4, 6, 7^ (Fig. 1a). Late on the morning of day 4, blastocysts are activated by a small estrogen surge that serves as the initiating signal for implantation^2, 8^. The blastocyst finally attaches to the luminal epithelium at midnight on day 4. The luminal epithelium initiates the formation of a uterine crypt after attachment, and the embryo can be observed at the bottom of the crypt^9^. Stimulation from embryo attachment seems to be transmitted from the endometrial epithelium to the stroma, where vascular permeability is increased (attachment reaction), and the surrounding stromal cells initiate differentiation into decidual cells (decidualization). By the evening of day 5, the endometrial luminal epithelium facing the attached blastocysts disappears, and trophoblast invasion is initiated.

**Fig. 1.**
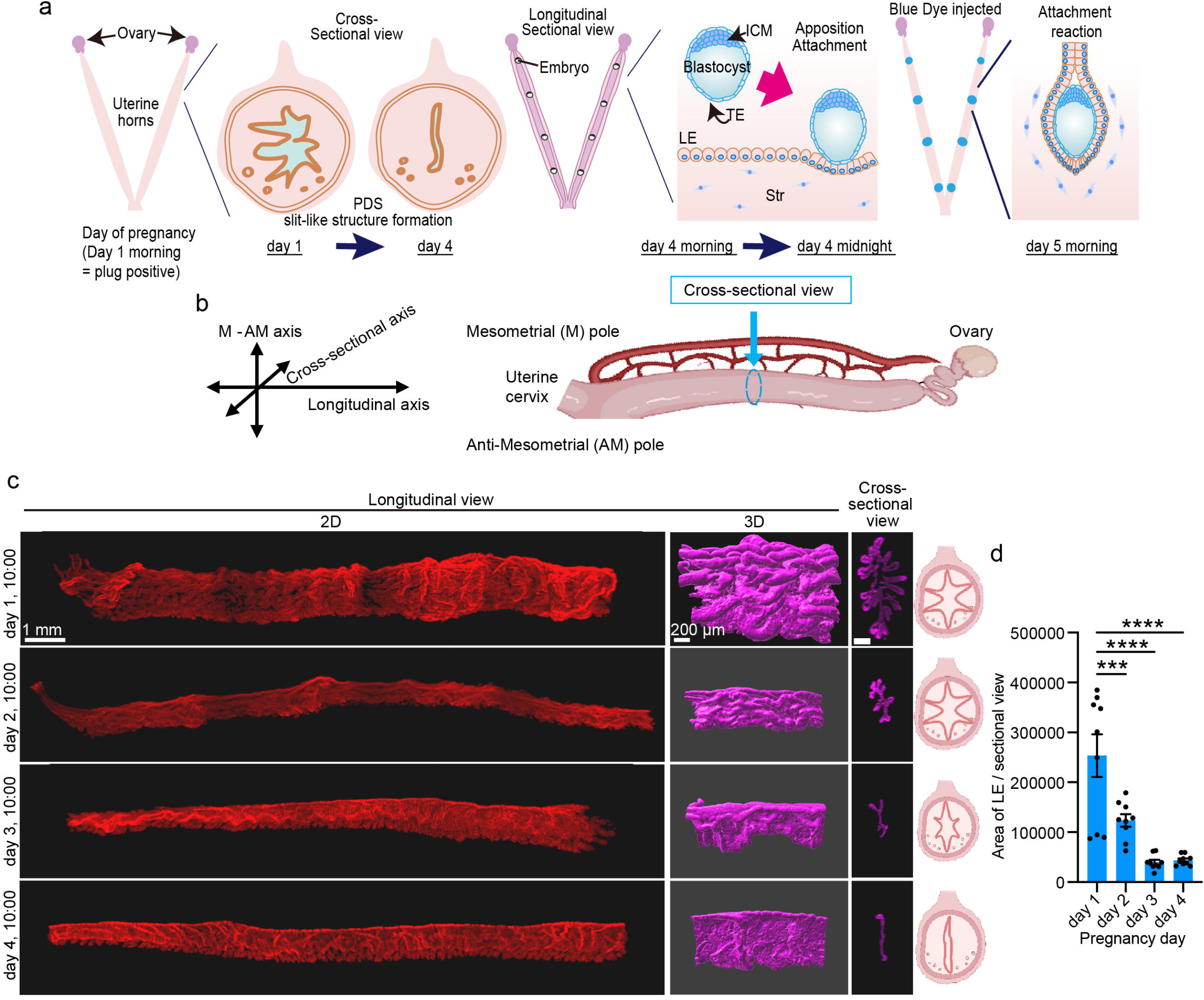
Three-dimensional observation of endometrial lumen morphology before embryo attachment. **a** A schematic diagram of embryo implantation. **b** Schematic diagrams of the longitudinal and cross-sectional views of the uterine horns. **c** Two-dimensional (2D) and three-dimensional (3D) views of the uterine epithelium stained for E-cadherin. The luminal epithelium was segmented and colored magenta using Imaris. Scale bar: 1 mm (left) and 200 µm (middle and right). Schematic diagrams of luminal shapes on each day of pregnancy are shown in the right panel. **d** The area of the luminal epithelium per cross section was quantified. Three sections were quantified per dam. The mean value of the three sections was used as a single biological replicate, and n = 3 dams were analyzed for each day of pregnancy. Data are presented as mean ± SEM, \*\*\**P* < 0.001, \*\*\*\**P* < 0.0001 by one-way ANOVA followed by Bonferroni’s post-hoc test.

In recent studies, our group and others have demonstrated that a series of luminal changes, namely PDS, formation of a slit-like luminal structure, and crypt formation, are crucial for embryo implantation^4, 5, 6, 7, 9, 10, 11^. For PDS and formation of the slit-like luminal structure, the P_4_-progesterone receptor (PGR) pathway is involved^4, 5, 6^. We previously reported that mice with epithelial deletion of PGR (*Pgr^fl/fl^;Ltf^Cre/+^* mice; *Pgr* eKO) showed sustained epithelial cell differentiation, resulting in defective uterine receptivity^6^. As for embryo attachment and subsequent crypt formation, the LIF-STAT3 pathway is crucial, in addition to P_4_-PGR signaling. Leukemia inhibitory factor (LIF) activates uterine LIF receptors (LIFRs) to evoke STAT3-mediated gene transcription, thus initiating the formation of implantation chambers (crypts)^9, 11, 12, 13, 14^. We have also reported that uterine-specific retinoblastoma knockout mice (*Rb* uKO) and enhancer of zeste homolog 2 knockout mice (*Ezh2* uKO) show impaired PDS because of cell cycle abnormalities in the endometrium^15, 16^. Notably, in these uKO models, although epithelial proliferation was continuously observed even on day 4, embryo attachment occurred, but subsequent embryo invasion was flawed. These differential phenotypes of uterine-specific gene deletions suggest that essential mechanisms other than PDS are involved in P_4_-and LIF-induced embryo attachment and subsequent pregnancy maintenance.

Accumulating evidence has demonstrated that PDS, slit-like formation, and crypt formation are important changes in the luminal epithelium for embryo attachment. However, how the overall changes in the luminal epithelium are dynamically regulated during the short period of embryo implantation remains unclear. It appears that PDS, slit formation, and crypt shaping occur sequentially; however, how each step influences the others remains uncertain. In particular, crypt formation occurs only in the presence of embryos^10^, indicating that epithelial changes require reciprocal interactions with embryos. To understand the molecular mechanism underlying these epithelial changes, we specifically examined p38α, a MAP kinase activated by phosphorylation in response to various environmental stresses and inflammatory cytokines, which regulates fundamental cellular processes such as proliferation, apoptosis, and cell differentiation^17, 18, 19^. Notably, p38α plays an important role in development and tissue differentiation by modulating the localization of E-cadherin, which is expressed in epithelial cells, thereby altering epithelial morphology^20, 21^. Embryo implantation requires cell differentiation and death, as well as glandular epithelial development and secretion, which may be influenced by p38α because it regulates various cell differentiation processes, including mammary gland duct formation^22, 23, 24^. Recently, deletion of p38α was reported to result in complete pregnancy failure in mice^25^, supporting our notion of a p38α-regulated mechanism shaping the endometrial epithelium during the peri-implantation period.

In this study, we first investigated the spatiotemporal changes in luminal shape in 3D spanning the period from just after coitus to the peri-attachment stage using a time-course analysis. Our analyses revealed that, prior to embryo attachment, the surface of the endometrial lumen not only became flattened but also developed distinct creases. The embryos were then attached to the flattened area, and crypts were formed. This epithelial shaping was impaired in mice with uterine-specific knockout (uKO) of p38α, resulting in failed embryo attachment. While treatment with P_4_ and LIF restored embryo attachment in p38α uKO mice, embryo invasion and subsequent pregnancy maintenance remained disturbed because of abnormal epithelial shaping, which was not rescued. In summary, we discovered that dynamic morphological changes in the endometrial lumen prior to implantation may influence the process of pregnancy establishment and maintenance, which is regulated by previously unknown mechanisms independent of P_4_-PGR and LIF-STAT3.

## RESULTS

### The morphology of the endometrial luminal surface dynamically changes before embryo attachment

To identify the spatiotemporal changes in the luminal epithelium prior to embryo attachment, we adopted 3D visualization, which recently revealed the detailed topography of the endometrial epithelium and the mechanism of embryo implantation^10^. Blood vessels enter the uterus from the mesometrium, orienting the uterus along the mesometrial–anti-mesometrial (M–AM) axis. Once blastocysts attach to the surrounding luminal cells, an implantation chamber (crypt) is formed by luminal epithelial (LE) evaginations toward the AM pole^10, 26, 27^ (Fig. 1a, b). Although the morphological changes in the endometrial lumen after embryo attachment have been reported^10, 28^, little is known regarding the luminal morphology prior to attachment. How the epithelial dynamics prior to embryo attachment influence subsequent pregnancy processes also remains unclear.

Therefore, we analyzed the luminal morphology in wild-type mice on the mornings (10:00) of days 1–4 of pregnancy. On day 1, the endometrial lumen extended in a disorderly manner against the M–AM plane. During days 2–4, the lumen became flattened, with some folding along the M–AM axis remaining (Fig. 1c). A cross-sectional view of the tissues revealed that the lumen became more slit-like, with the epithelial mass decreasing daily prior to embryo attachment (Fig. 1c, d), which was consistent with the luminal changes previously depicted using conventional histology^4, 29^ (Supplementary Fig. 1a).

Next, we investigated the relationship between luminal folding and blastocyst positioning by observing day 4 endometria from the morning to midnight, when blastocysts arrived in and attached to the uterine lumen, respectively. We observed the 3D morphology of the uterine lumen and the positions of the blastocysts on day 4 morning (day 4 10:00), day 4 evening (day 4 16:00), day 4 night (day 4 20:00), day 4 midnight (day 5 00:00), and day 5 morning (day 5 10:00). Once embryo attachment occurred, vascular permeability increased in the surrounding endometrium, which could be observed by injecting blue dye into the mice (blue dye reaction)^30^. The blue dye reaction rates were 0% (0/5) at 16:00 on day 4, 18.8% (3/16) at 20:00 on day 4, 66.7% (4/6) at 00:00 on day 5, and 100% (7/7) at 10:00 on day 5 (Fig. 2a–c). Embryo attachment was completed in more than half of the animals by midnight on day 4; therefore, we defined the evaluation time immediately before embryo attachment as day 4 20:00. As previously described, the surface of the endometrial lumen was flattened at day 4 10:00. On day 4, from 16:00 to 20:00, the flat lumen exhibited alternating shrunken and stretched areas. The shrunken and stretched regions were clearly distinguished. Shrunken areas with multiple folds were observed at regular intervals. No regularity in the positions of the embryos was observed until day 4 16:00; however, by day 4 20:00, the embryos were observed in the shrunken areas, and were increasingly localized to the stretched areas at later time points. At day 4 midnight, the embryos were attached to the AM pole of the lumen in the stretched areas, and crypts were then formed on the morning of day 5 (day 5 10:00) (Fig. 2d). Notably, this luminal morphology was comparable between conventional two-dimensional section analysis and three-dimensional evaluation (Supplementary Fig. 1b). To quantify the transition between shrunken and stretched areas in wild-type mice before and after embryo attachment, the angles of the luminal foldings against the M–AM axis were quantified, as previously reported^31^, showing that shrunken areas have vertical folding (around 0 degree), whereas stretched areas have horizontal folding (around 90 degree) on day 4 20:00 (Fig. 2e). The IS area showed smaller folding angles than the inter-IS area, consistent with a greater proportion of longitudinal folds, reflecting the difference between the shrunken and stretched areas. To investigate the relationship between embryo location and luminal folds, the angles of the luminal epithelia against the longitudinal axis were additionally measured (the embryo center and ±30 μm from the center). This analysis revealed a significant difference between day 4 20:00 (pre-attachment) and day 5 00:00 (post-attachment) (Fig. 2f), indicating that embryo location differed between the pre-and post-attachment time points. These results demonstrate that the uterus, just prior to embryo attachment, underwent dynamic changes in luminal morphology once the embryos arrived.

**Fig. 2.**
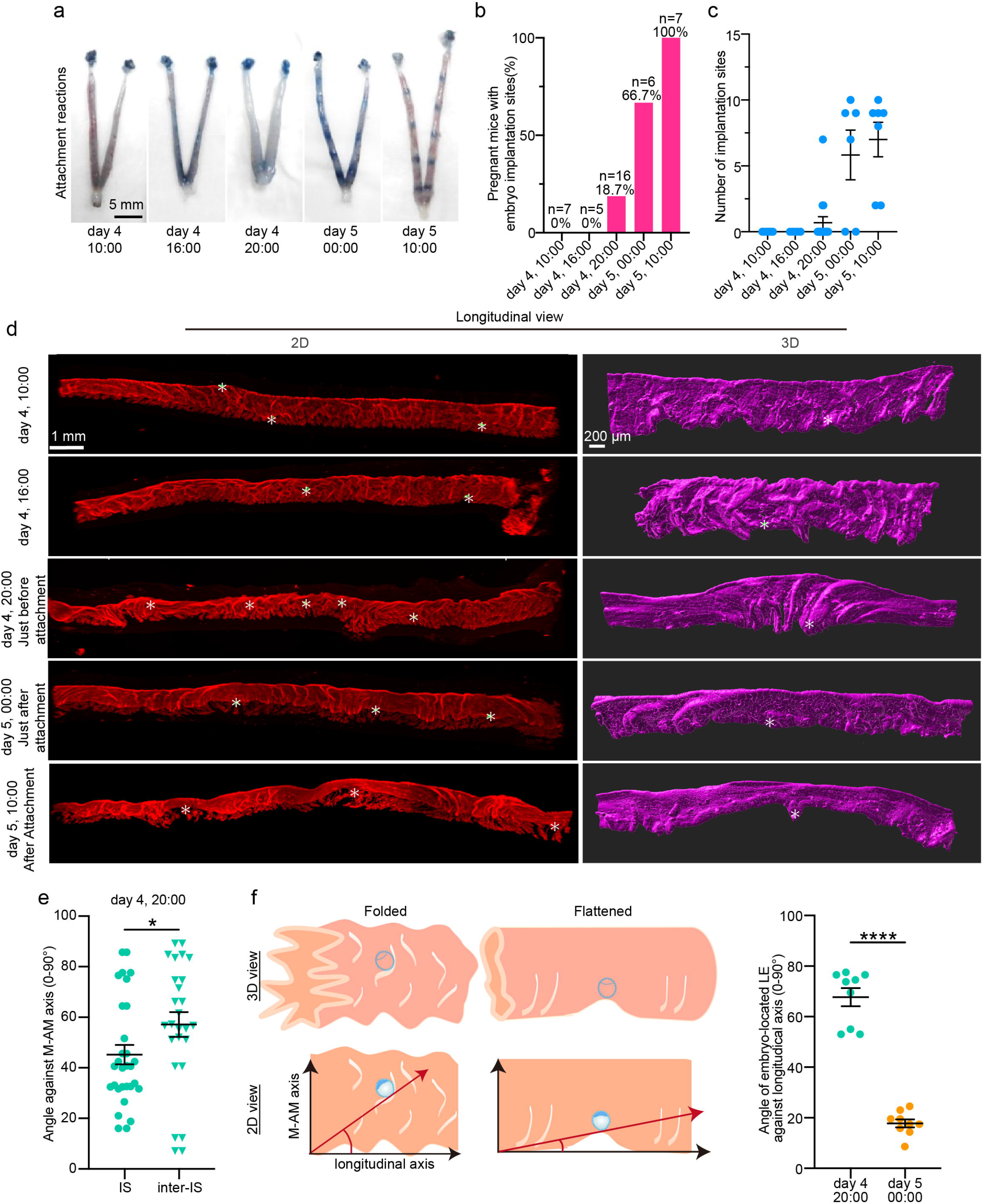
Dynamic morphological changes in the endometrial luminal epithelium occur on the night of day 4, just before embryo attachment. **a** Representative images of pregnant uteri from the morning of day 4 morning to the morning of day 5 following blue dye injection to depict embryo attachment sites. Scale bar: 1 cm. **b** Percentage of implantation-positive females in (**a**). The number of replicates and the percentage of implantation-positive females are shown above each bar. **c** The number of implantation sites in (**a**) and (**b**). Data are presented as mean ± SEM. **d** 2D and 3D longitudinal views of uterine epithelia stained for E-cadherin. In the right panels, the luminal epithelium is segmented and colored magenta using Imaris. Scale bar: 1 mm (left) and 200 µm (right). Asterisks indicate the locations of embryos. **e** Angle of luminal folding against M–AM axis on day 4 at 20:00 around the implantation sites (IS) and inter-implantation sites (inter IS). n = 4 dams for each time point of pregnancy were quantified. **f** Angle of the luminal epithelium relative to the longitudinal axis at three positions on day 4 20:00 and day 5 00:00. Angles were measured at three positions for each sample, and their mean was used for quantitative analysis. n = 3 dams per time point.

### Uterine p38α activation is crucial for luminal morphology and subsequent embryo attachment

We then examined how luminal dynamics before attachment were regulated. We performed single-cell RNA-seq (scRNA-seq) analysis for day 4 and day 5 uteri at 10:00 (Fig. 3). Uterine tissues contain multiple cell types, including epithelial, stromal, vascular endothelial (VE), and immune cells (Fig. 3a, Supplementary Fig. 2a, and Tables S1 and S2). Among them, luminal cell clusters (LE) can be divided into two types: conventional luminal cells (LE), which are predominantly observed on day 4, and activated LE (LE_activated), whose population increases on day 5 (Fig. 3a). To determine the characteristics of the LE_activated cells, we performed enrichment analyses using Enrichr^32^ (Fig. 3b, Table S2). Transcription factor protein–protein interactions (PPIs) revealed enrichment of ATF2 and JUNB, which are related to stress signalling^33^. In agreement with this result, pathway analysis using MSigDB Hallmark also revealed TNFα-related and hypoxia signaling. Furthermore, gene ontology analysis using Metascape^34^ identified pathways related to oxidative stress and cell motility (Fig. 3c, Table S2). We then investigated stromal cell types in the same manner (Fig. 3d, Supplementary Fig. 2b, and Table S3). These were clustered into five types: non-proliferative (Non-pro), proliferative (Pro), sub-epithelial (Sub-epi), sub-endothelial (Sub-endo), and attached (Attached). As the day 5 stroma exclusively contained the Attached cluster, we analyzed the enriched pathways in this cell type. Similar to LE_ activated, this cluster was enriched in stress-related transcription factors, signaling pathways, and Gene Ontology (GO) terms (Fig. 3e, f, and Table S4).

**Fig. 3.**
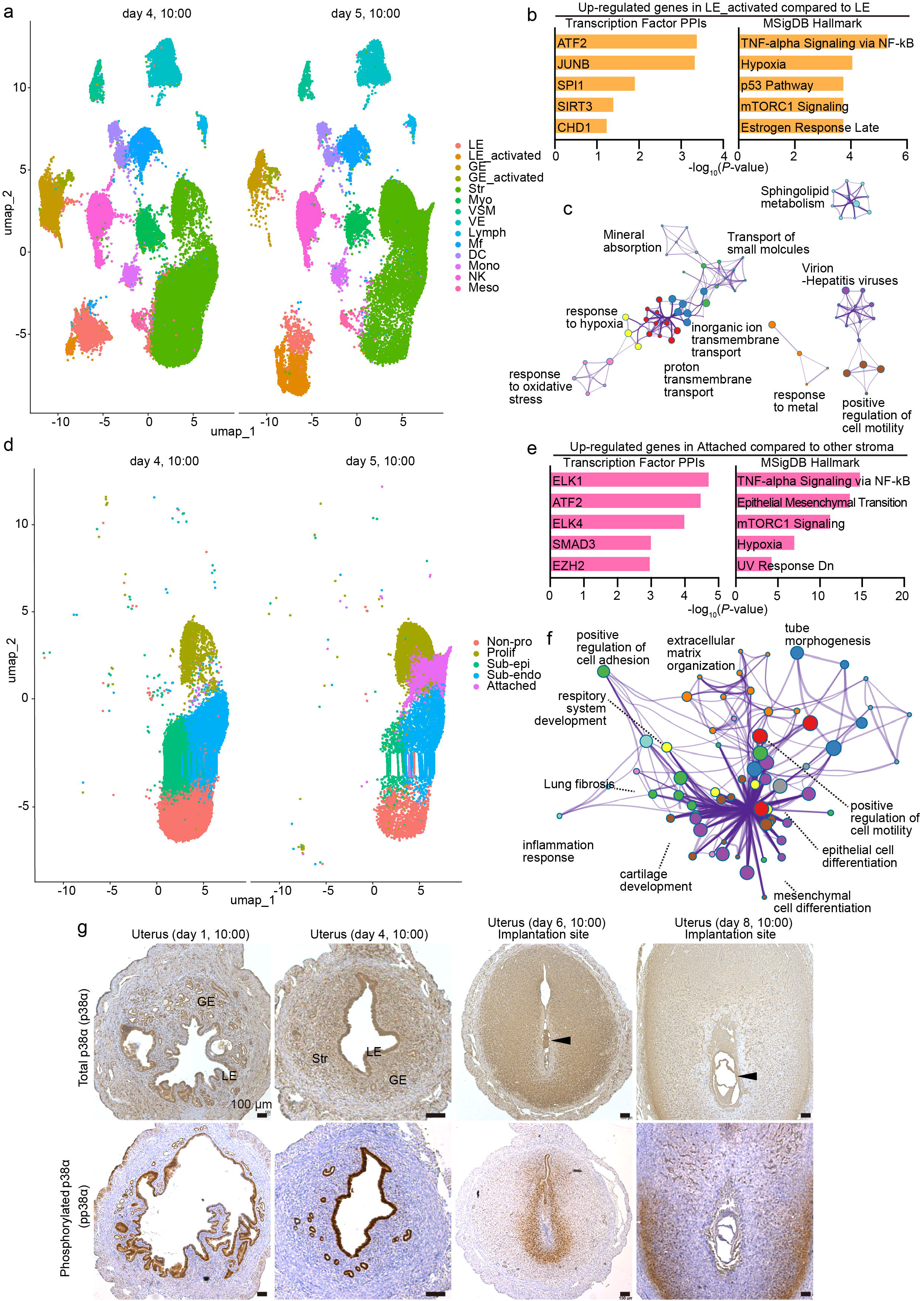
Stress-related signals are activated in the uterus during the peri-attachment phase. **a** UMAP of scRNA-seq cell types on days 4 and 5 of pregnancy. Dots indicate individual cells, and colors indicate different clusters. **b** Enrichment analyses for upstream transcription factors (left) and Gene Ontology (GO) (right) of highly expressed genes in the LE_activated vs. LE cells. **c** Network of GO terms related to the upregulated genes in the LE_activated cells compared with those in LE cells. Each node represents an enriched term and is colored according to its cluster ID. **d** UMAP of different stromal cell types on days 4 and 5 of pregnancy. Dots indicate individual cells, and colors represent different clusters. **e** Enrichment analyses for upstream transcription factors (left) and Gene Ontology (GO) (right) of highly expressed genes in the Attached cluster compared with those in the other stromal clusters. **f** Network of GO terms related to the upregulated genes in the Attached cells compared with those in the other stromal clusters. Each node represents an enriched term and is colored according to its cluster ID. **g** Representative images of p38α (top) and phosphorylated p38α (pp38α; bottom) immunohistochemistry during days 1, 4, 6, and 8 of pregnancy. Scale bar: 100 µm. LE: luminal epithelia, GE: glandular epithelia, Str: Stroma, Em; embryo, Le; Luminal epithelium, St; Stroma. Arrowheads indicate embryos. At least three independent dams were evaluated for each day of pregnancy.

As a possible regulator of day 5-specific LE and stromal cell types, we focused on p38α, a MAP kinase protein. p38α is phosphorylated for activation in response to various kinds of cellular stimuli^18^. Notably, phosphorylated p38α (pp38α) translocates into the nuclei to activate ATF2-JUN-induced transcription of cytokines, including TNFα^33^. Our immunostaining experiment revealed that p38α was highly expressed and activated by phosphorylation in the pre-attachment luminal epithelia (Fig. 3g), indicating a role for p38α in the luminal epithelium before embryo attachment. After attachment, pp38α was expressed in both the endometrial epithelium and stroma, especially around the attached embryos on days 6 and 8.

To investigate the roles of uterine p38α, we established mice with uterine-specific deletion of p38α (p38α uKO) by crossing p38α-floxed mice with those carrying a *Pgr*-Cre driver. Efficient deletion and inactivation of p38α protein in p38α uKO mice was confirmed by immunostaining for p38α and pp38α (Fig. 4a). To examine the reproductive phenotypes of p38α uKO and littermate p38α-floxed females (p38α Ctrl), we mated them with fertile wild-type (WT) male mice. p38α uKO mice showed complete infertility (Fig. 4b). As we detected comparable numbers of hatched blastocysts by flushing the uterine cavity with saline solution on the morning of day 4 in each genotype (Fig. 4c), uterine p38α did not influence embryo development before attachment. We then intravenously injected Chicago blue dye solution to examine the implantation sites on day 5 of pregnancy. However, p38α uKO uteri showed no attachment sites on day 5 of pregnancy, and blastocysts were recovered by saline flushing of the uterine cavities (Fig. 4d), indicating failed embryo attachment. On day 6 of pregnancy, the number of embryo attachment sites was significantly reduced in p38α uKO mice compared with that in the p38α uCtrl mice (Fig. 4e). Furthermore, our 3D imaging on the morning of day 5 revealed failed crypt formation in p38α uKO mice, suggesting that p38α uKO mice are infertile because of embryo attachment failure (Fig. 4f).

**Fig. 4.**
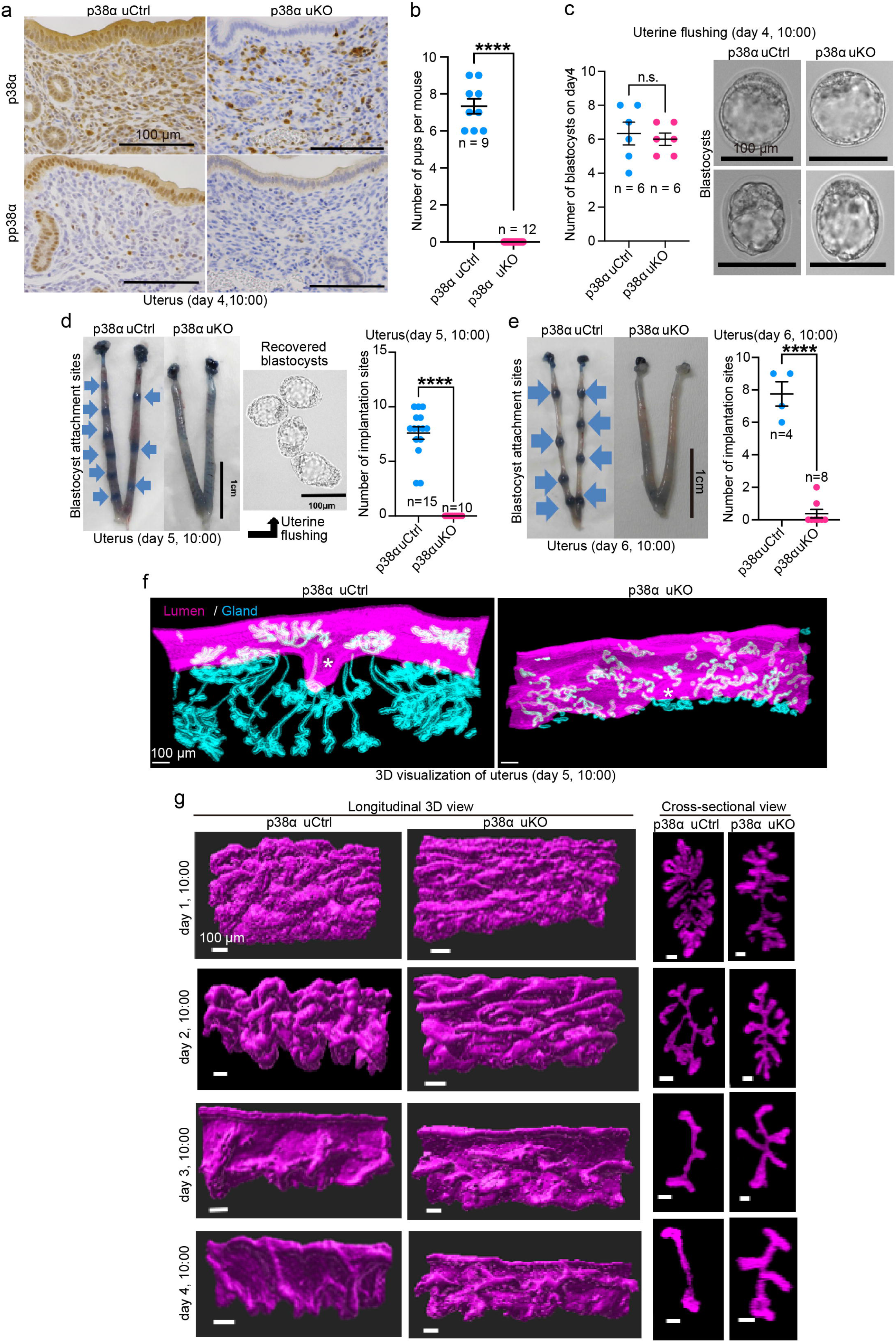
Morphological changes in the pre-attachment endometrial luminal epithelium are impaired in uterus-specific p38α KO mice. **a** Efficient p38α deletion was confirmed by immunostaining for p38α and phosphorylated p38α (pp38α) in the uteri on day 4 of pregnancy. Scale bar = 100 µm. **b** Average litter size for each genotype. The numbers of replicates are shown in the graph. Data represent the mean ± SEM, and \*\*\*\**P* < 0.0001 by Student’s t-test. **c** Comparable numbers of flushed blastocysts (left) and their morphology (right). The graph shows the number of dams tested. Data represent the mean ± SEM, n.s.: not significant according to Student’s t-test. Scale bar = 100 µm. **d** Representative images of the uteri from each genotype (left) and the average number of implantation sites on day 5 of pregnancy. Flushed embryos from p38α uKO are shown next to the uterine images. Scale bar = 1 cm (uteri) and 100 µm (embryos). The number of replicates is shown on the graph. Data represent the mean ± SEM, and \*\*\*\**P* < 0.0001 by Student’s *t*-test. **e** Representative images of the uteri from each genotype (left) and the average number of implantation sites on day 6 of pregnancy. Scale bar = 1 cm. The number of replicates is shown on the graph. Data represent the mean ± SEM, and \*\*\*\**P* < 0.0001 by Student’s t-test. **f** Representative 3D images of day 5 pregnant uteri stained for E-cadherin. The luminal and glandular epithelia were segmented and colored magenta and cyan, respectively. Scale bar = 100 µm. n=4 dams were evaluated for each group. **g** Representative 3D views of luminal epithelia during days 1–4 of pregnancy, segmented based on epithelial staining for E-cadherin. Scale bar = 100 µm. n=3 dams were evaluated for each group.

Our observations indicating luminal activation of p38α during the pre-attachment period (Fig. 3g), as well as infertility with failed embryo attachment in p38α uKO females (Fig. 4), motivated us to investigate the involvement of this molecule in luminal dynamics. We then compared the morphological changes in the lumen of p38α uCtrl and p38α uKO mice from days 1 to 4 using 3D imaging (Fig. 4g). Similar to the observations in wild-type mice, p38α uCtrl mice showed that folding of the endometrial lumen along the M–AM axis gradually disappeared, although some folding remained. Accordingly, the endometrial lumen flattened as pregnancy progressed from days 1 to 4. In contrast, in p38α uKO mice, folding along both the cross-sectional and longitudinal axes remained even on day 4, which appeared as serrated luminal shapes in the 2D view. Considering that embryo attachment failed in p38α uKO mice, the luminal changes during the pre-attachment phase could influence successful embryo attachment.

### Supplementation with P_4_ and LIF, two major factors supporting embryo implantation, rescues flawed embryo attachment, but not subsequent pregnancy maintenance, in p38α uKO mice

Our group and others have previously revealed that PDS, wherein epithelial cell proliferation is terminated before implantation, is an indicator of endometrial embryo receptivity^5^. P**_4_** is an inducer of PDS as well as slit formation in the endometrial lumen^4, 6^. Immunostaining for Ki67, a cell proliferation marker, showed an increased number of proliferating cells in the luminal epithelium of p38α uKO mice on the morning of day 4, suggesting that PDS was impaired in this milieu (Fig. 5a, b). This result motivated us to examine whether P**_4_** supplementation could rescue the phenotype of p38α uKO. We thus treated p38α uKO mice with P**_4_** during the preimplantation period and examined the resulting morphological changes in the endometrial lumen using 3D analysis. On days 3 and 4, P_4_ treatment suppressed folding in both the cross-sectional and longitudinal axes, thus flattening the lumen (Fig. 5c–e). Furthermore, PDS was rescued by P_4_ injection in p38α uKO mice (Fig. 5f, g). Notably, p38α uKO showed normal hormone-producing capacity of the ovary, as serum estradiol-17β (E₂) and progesterone (P_4_) levels were comparable (Supplementary Fig. 3a). We also confirmed that the expression of estrogen receptor (ERα) and progesterone receptor (PGR) in the uterus was not influenced by p38α deletion (Supplementary Fig. 3b). Although luminal morphology was improved by P_4_ treatment, embryo attachment still failed in the mutant mice (Fig. 5h), suggesting that some other factor is required for p38α-dependent embryo attachment.

**Fig. 5.**
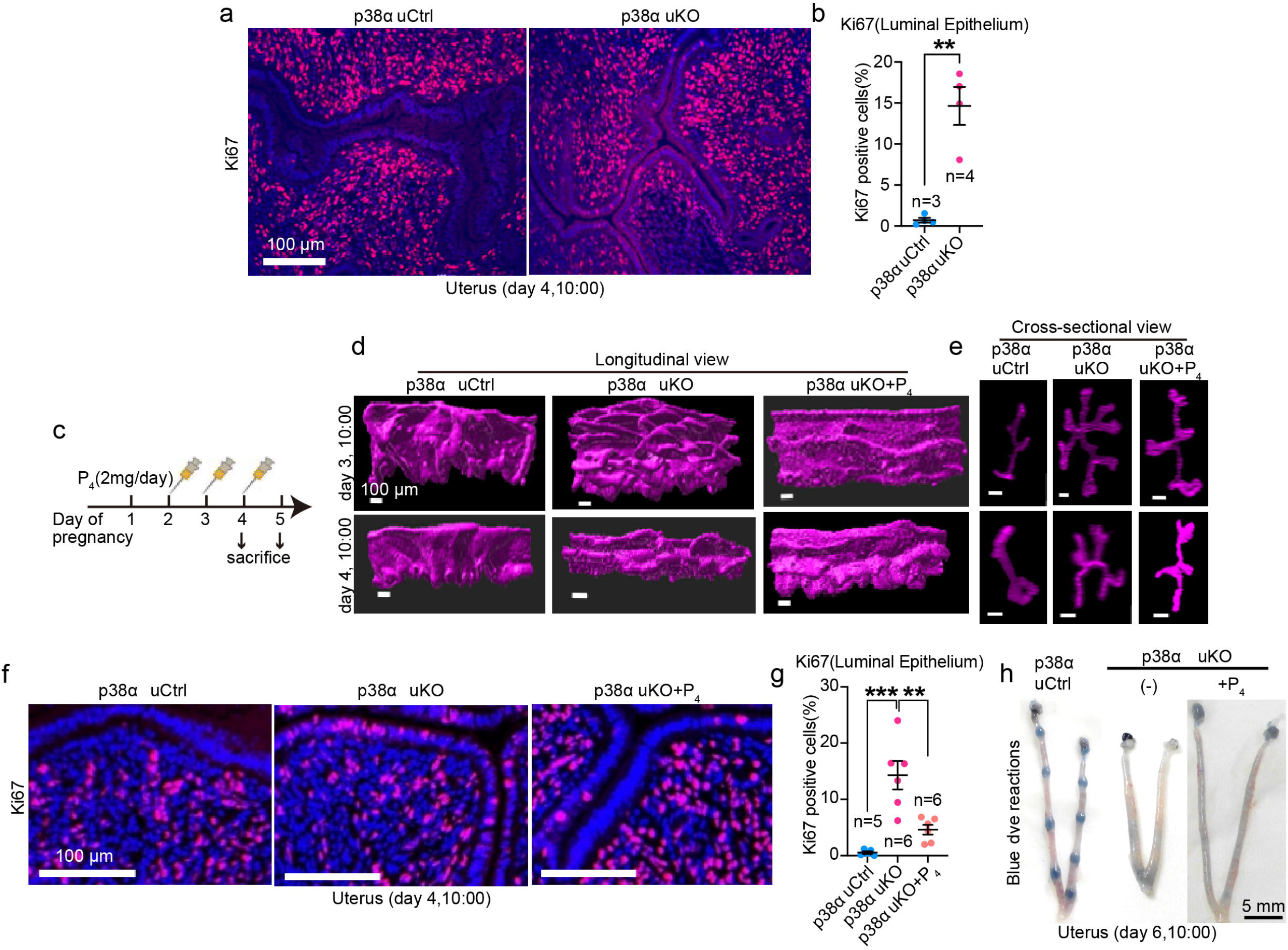
P_4_ supplementation to p38α uKO mice partially rescues structural changes and proliferation–differentiation switching (PDS) in the endometrial luminal epithelium before embryo attachment. a, **b** Representative images of Ki67 immunofluorescence in the luminal epithelium of the uteri on the morning of day 4. The percentages of Ki67-positive cells among total luminal cells are shown in (**b**). The number of replicates is shown on the graph. Data represent the mean ± SEM, \*\**P* < 0.01 by Student’s *t*-test. Scale bar = 100 µm. **c** Schedule of P_4_ treatment in p38α uKO females during days 1 to 5 of pregnancy. **d** Representative 3D longitudinal views of the luminal epithelia from control and p38α uKO uteri with or without P_4_ treatment, segmented from epithelial staining for E-cadherin. Scale bar = 100 µm. **e** Cross-sectional view of the luminal epithelia in (**d**). Scale bar = 100 µm. n=3 dams were evaluated for each group in (d) and (e). **f, g** Representative images of Ki67 immunofluorescence in the luminal epithelium on the morning of day 4 in uteri from control and p38α uKO mice with or without P_4_ treatment. The percentages of Ki67-positive cells per total luminal cells are shown in (**g**). The number of replicates is shown on the graph. Data represent the mean ± SEM, \*\**P* < 0.01 and \*\*\**P* < 0.001 by one-way ANOVA followed by Bonferroni’s post-hoc test. Scale bar = 100 µm. **h** Representative images of the uteri from control and p38α uKO mice with or without P_4_treatment on day 6 of pregnancy. Scale bar = 5 mm.

Besides the P_4_-PGR pathway, leukemia inhibitory factor (LIF) is a critical factor for embryo attachment^9, 11, 35^. LIF is an interleukin-6 family member secreted by the endometrial glands on the morning of day 4^35, 36, 37^. *In situ* hybridization showed significantly decreased LIF expression in the p38α uKO uterus on the morning of day 4 even after P_4_ treatment (Fig. 6a). Furthermore, activation of STAT3, a transcription factor downstream of LIF, was also downregulated, as determined by immunostaining for phosphorylated STAT3 in the p38α uKO uterus on the morning of day 4 (Fig. 6b).

**Fig. 6.**
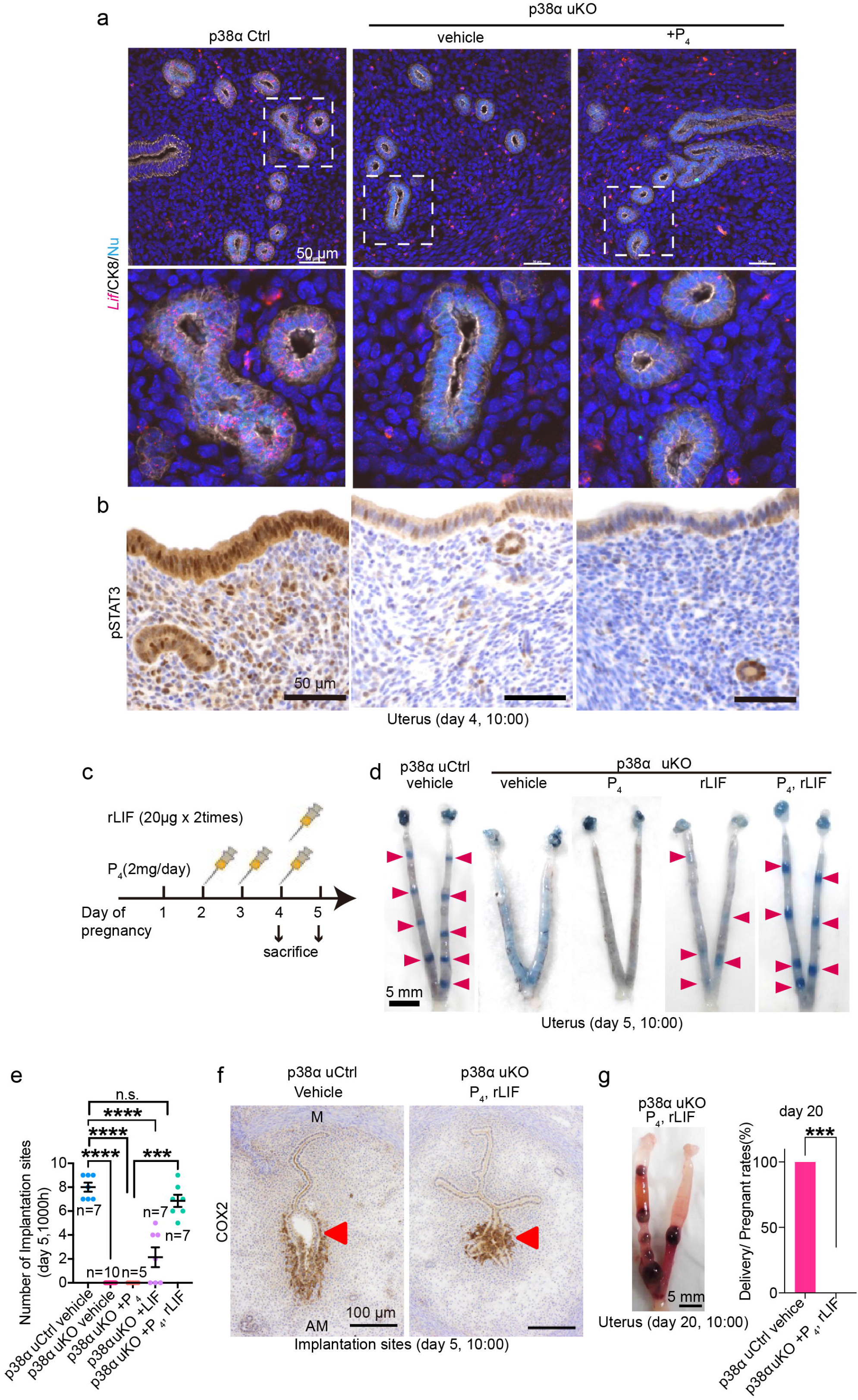
Impaired LIF-STAT3 signaling affects embryo attachment in p38α uKO mice. **a** Representative images of *Lif in situ* hybridization (top and middle) and phosphorylated STAT3 (pSTAT3) immunostaining (bottom) in day 4 uteri from control and p38α uKO mice with or without P_4_ treatment. Epithelial cells immunostained for CK-8 are shown in the top and middle panels. The area indicated by a dashed line in the top panel is shown in the middle panel. Scale bar = 50 µm. **b** Schedule of P_4_ and rLIF treatment in p38α uKO female mice during days 1 to 5 of pregnancy. n=3 dams were evaluated for each group in (a) and (b). **c** Protocol for P₄ and LIF treatment. **d** Representative images of day 5 pregnant uteri from control and p38α uKO mice with or without P_4_ and rLIF treatment. Arrowheads indicate sites of embryo attachment. Scale bar = 5 mm. **e** The number of implantation sites in (**d**) was calculated. The number of replicates is shown on the graph. Data represent the mean ± SEM, \*\*\**P* < 0.001, \*\*\*\**P* < 0.0001, and n.s.: not significant by one-way ANOVA followed by Bonferroni’s post-hoc test. **f** Representative images of COX2 immunostaining at day 5 implantation sites from control and p38α uKO uteri treated with P_4_ and rLIF. Scale bar = 100 µm. M: mesometrial pole; AM: anti-mesometrial pole. Arrowheads indicate embryos. **g** Representative images of day 20 pregnant uteri from p38α uKO mice treated with P_4_ and rLIF. The percentage of deliveries among the total number of pregnant females is shown in the graph on the right. The number of replicates is shown on the graph. Data represent the mean ± SEM, \*\*\**P* < 0.001 by Student’s *t*-test.

Based on these findings, we investigated whether supplementation with recombinant LIF (rLIF) in combination with P**_4_** could rescue the defective embryo attachment observed in p38α uKO mice. Five groups were established: vehicle administration to p38α uCtrl and p38α uKO mice, P**_4_** alone to p38α uKO mice (at 10:00 on days 2–4), rLIF alone to p38α uKO mice (at 9:00 and 18:00 on day 4), and both P**_4_** and rLIF to p38α uKO mice (Fig. 6c). The number of embryo attachment sites following treatment with either P**_4_** or rLIF alone did not differ from that in the vehicle group. In contrast, simultaneous supplementation with P**_4_** and rLIF increased the number of embryo attachment sites in p38α uKO mice to a level comparable to that in the p38α uCtrl group, as shown by the clear blue dye reactions on the morning of day 5, indicating that both P**_4_** and LIF are required for p38α-dependent embryo attachment (Fig. 6d, e). COX2 (Fig. 6f) and phosphorylated STAT3 (Supplementary Fig. 4), which are induced at embryo attachment sites^11^, were expressed in p38α uKO mice upon P**_4_** and rLIF treatment, further supporting our notion. However, despite embryo attachment, P_4_ and rLIF treatment could not restore full-term pregnancy with litters in p38α uKO mice (Fig. 6g). On day 8, although the number of implantation sites was comparable between the two groups, none of the implantation sites underwent normal embryogenesis, and hematomas formed in p38α uKO treated with P**_4_** and rLIF (Supplementary Fig. 5), indicating that pregnancy was not maintained normally. Eventually, P_4_-and rLIF-treated p38α uKO females failed to give birth and exhibited severe embryo resorption (Fig. 6g). This suggests that p38α is required for P_4_-and LIF-induced embryo attachment as well as healthy pregnancy maintenance.

### p38α plays an important role in establishing luminal epithelial morphology for embryo attachment, and the changes by the morning of day 4 provide an important scaffold for embryo attachment site formation

Flawed pregnancy maintenance in the mutant, even after treatment, led us to examine how pregnancy events after embryo attachment were disturbed in p38α-deleted uteri. We thus investigated the luminal morphologies on days 5 and 6, when embryo attachment and invasion were evident (Fig. 7a). We found that folding was evident along the longitudinal axis in p38α uKO regardless of any treatment. Simultaneous treatment with P_4_ and rLIF created a crypt on day 6, but longitudinal folding was still observed around the crypt. Furthermore, crypt formation was poorly initiated on the morning of day 5 in P_4_ and rLIF-treated p38α uKO mice, showing obvious longitudinal folding. Considering that this folding should be eliminated by the night of day 4 in normal pregnancy (Fig. 2), we investigated the luminal morphology at day 4 20:00, just before embryo attachment (Fig. 7b). Similar to the preparatory phase of embryo attachment on the morning of day 4 (10:00 am), the surface of the stretched lumen, i.e., the site of embryo attachment, was flat in p38α uCtrl. In contrast, p38α uKO and p38α uKO+ rLIF (P_4_ non-treated group) showed remarkable persistence of luminal folding in the longitudinal axis with a poorly established stretched area. In p38α uKO+P_4_ and p38α uKO+ rLIF +P_4_ (P_4_ treated group,) flattening and stretching of the lumen was partially rescued, but longitudinal folding remained (Fig. 7b, left). We also observed the luminal shapes in cross-sectional views to examine the relationship between the shape of the slit-like lumen and embryo location (Fig. 7b, right, c, and d). In the p38α uCtrl, the stretched luminal areas serving as embryo attachment sites, were flat and slit-like, and embryos were preferentially positioned toward the AM pole (Fig. 7b, right). In contrast, regardless of treatment, failure of luminal flattening in p38α uKO mice was accompanied by less stereotyped embryo positioning. These abnormal morphologies of the luminal epithelium appeared as increased epithelial mass (Fig. 7c) and increased epithelial branching in the uKO (Fig. 7d). As in wild-type mice, the angles of luminal foldings against the M–AM axis on day 4 at 20:00 were also evaluated in both the IS and inter-IS regions of p38α uKO mice. In contrast to uCtrl, no significant difference was observed between the IS and inter-IS regions in the uKO uteri (Fig. 7e). Overall, these results suggest that p38α is responsible for morphological changes in the lumen before embryo attachment, which P_4_, but not LIF, partially assists. Considering that P_4_ could not solely restore embryo attachment in p38α uKO, LIF seems to act as an inducer of attachment under the influence of P_4_.

**Fig. 7.**
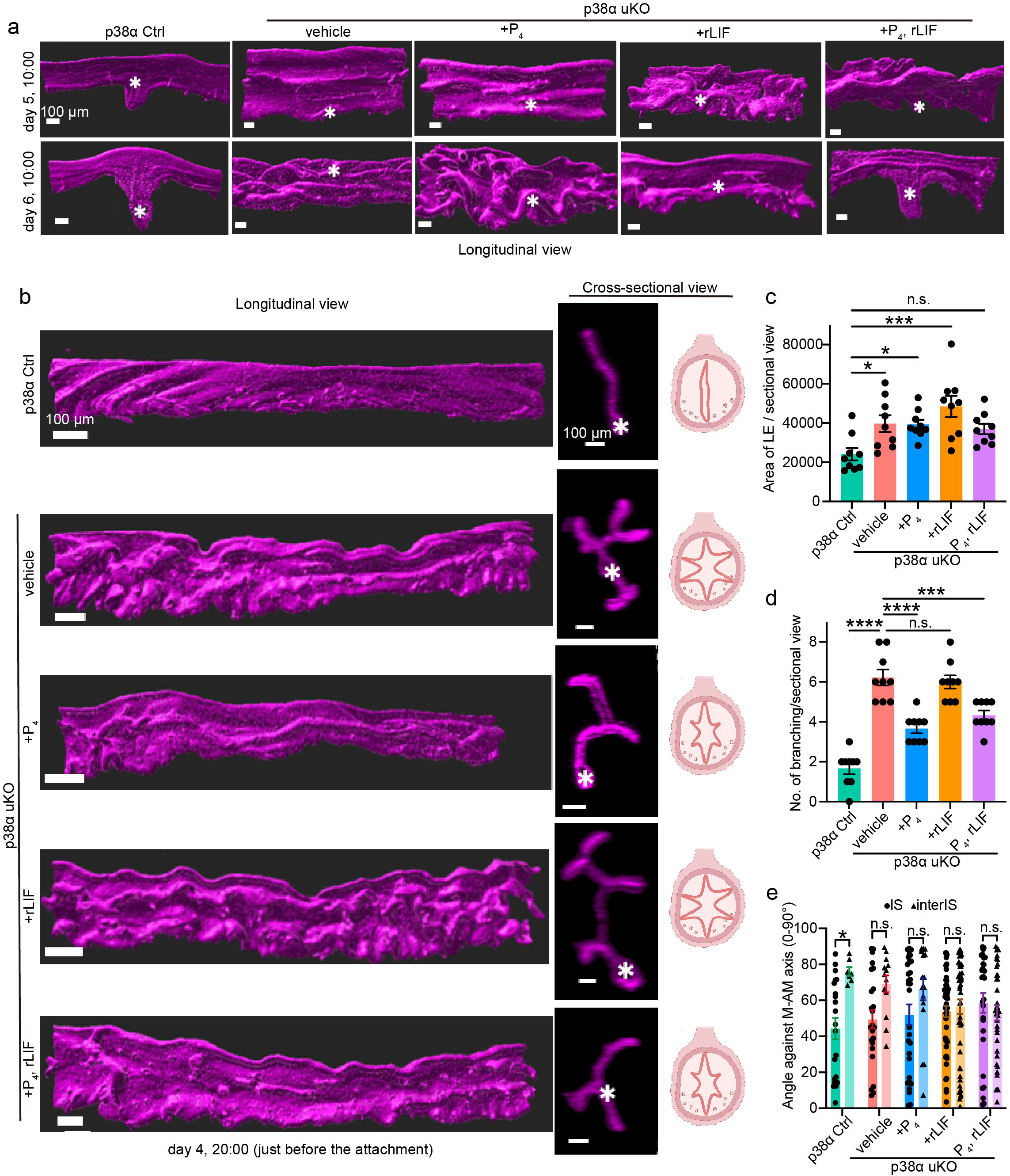
Embryo attachment in p38α uKO mice is rescued by supplementation with P_4_ and LIF, but luminal positioning remains inappropriate. **a** Representative 3D views of the luminal epithelia collected from each genotype on days 5 and 6 of pregnancy. Scale bar = 100 µm. Asterisks indicate embryos. **b** Representative 3D views of the luminal epithelium on day 4 midnight, immediately before embryo attachment, collected from each genotype. Scale bar = 100 µm, asterisks indicate embryos. The graphs in the right panels show the luminal shapes for each genotype and condition. **c** Average area per cross section calculated from the images shown in (**b**). Data represent the mean ± SEM, and *P*-values were determined by one-way ANOVA followed by Bonferroni’s post hoc test. **d** Average number of luminal branches per cross section calculated from the images shown in (**b**). Three cross-sectional images were quantified per dam, and the mean of the three measurements was used as one biological replicate (n = 3 dams per group). **e** Angle of luminal folding against the M–AM axis on day 4 at 20:00. n = 3 (uKO+vehicle), 4 (uCtrl, uKO+rLIF, uKO+P_4_, rLIF), 5 (uKO+P_4_) dams were quantified. Data represent the mean ± SEM, and \**P* < 0.05, \*\*\**P* < 0.001, \*\*\*\**P* < 0.0001, n.s.: not significant by one-way ANOVA followed by Bonferroni’s post-hoc test.

### Uterine p38α supports epithelial–stromal communication during embryo attachment

We then examined how p38α influences feto-maternal interactions to accomplish healthy embryo implantation and luminal dynamics.

As *Pgr*-Cre mediates recombination in both the epithelial and stromal compartments of the uterus, we next examined whether epithelial loss of p38α alone could account for the severe implantation phenotype observed in p38α uKO mice. To this end, we generated epithelial-specific p38α knockout (eKO) mice using the *Ltf*-iCre driver. In contrast to *Pgr*-Cre-mediated p38α uKO females, which showed complete infertility and severe defects in luminal remodeling, *Ltf*-iCre-mediated epithelial-specific deletion of p38α did not critically impair female fertility and resulted only in mild uterine abnormalities (Supplementary Fig. 6). These results indicate that epithelial p38α loss alone is insufficient to reproduce the severe phenotype of p38α uKO mice. This finding prompted us to examine whether p38α regulates luminal remodeling through epithelial–stromal communication rather than through an exclusively epithelial-autonomous mechanism. We therefore performed scRNA-seq in control uteri on day 4 at 20:00 and day 5 00:00, and in p38α uKO uteri on day 5 at 00:00 with or without P_4_+ rLIF treatment. (Fig. 8a, Table S5). Similar to Fig. 3a, we found multiple cell types in the uteri, including epithelial, stromal, myometrial, and immune cells. Among these, we focused on stromal cells, as their population increased during the embryo attachment process in the uCtrl (Fig. 8a). The stromal cells were further clustered into non-proliferative (Non-prolif), proliferative (Pro), decidualizing (Dec), sub-epithelial (Sub-epi), sub-endothelial (Sub-endo), and uKO-specific clusters (Fig. 8b, Table S6). The uKO-specific cluster was highly enriched in uKO uteri. We then traced stromal differentiation using pseudo-time analysis (Fig. 8c and d). Intriguingly, the uKO-specific cluster was less differentiated than the other stromal cell types, except for the Non-prolif cluster (Fig. 8d).

**Fig. 8.**
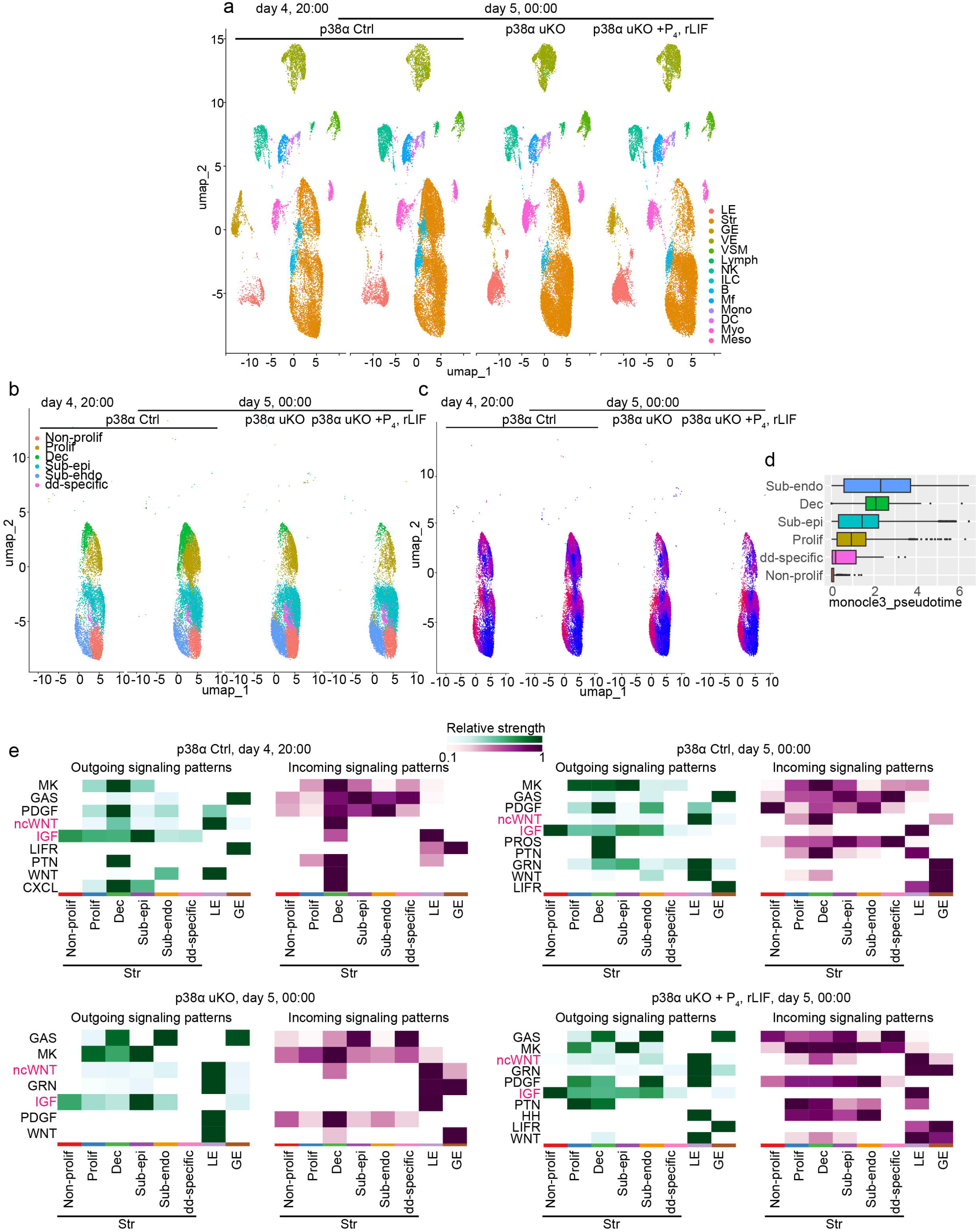
p38α plays an important role in stromal differentiation to ensure appropriate epithelial–stromal interactions before embryo attachment. **a** UMAP of scRNA-seq showing various cell types from control uteri on day 4 evening and midnight, and from p38α uKO with or without P_4_ and rLIF treatment on day 4 midnight. Dots indicate individual cells, and colors indicate different clusters. **b** UMAP of scRNA-seq showing stromal cell types from control uteri on day 4 evening and midnight, and from p38α uKO with or without P_4_ and rLIF treatment on day 4 midnight. Dots indicate individual cells, and colors indicate different clusters. **c** UMAP plots show the diffusion pseudotime of stromal cells. The colors of the dots indicate the pseudo-time from early (blue) to late (dark red). **d** Boxplot showing the distribution of pseudo-time within different stromal cell clusters. Colors and labels indicate the cell types corresponding to those shown in (**b**). **e** Heatmap showing the outgoing (left) or incoming (right) communication strengths of key pathways between the major uterine endometrial cell types in each genotype and condition.

We then examined the signal transduction between the epithelial (LE and GE) and stromal clusters, focusing on secretory molecules that can bridge the epithelial and stromal compartments (Fig. 8e). In uCtrl, non-canonical WNT (ncWNT) from the LE and stroma and IGF from the stroma affected the decidualizing and luminal cells, which were enhanced upon embryo attachment (Fig. 8e, upper). In contrast, in p38α-deficient uteri, only epithelial cells strongly sent and received signals (Fig. 8e, lower left), which remained even with P_4_ and rLIF treatment (Fig. 8e, lower right). These results demonstrate flawed epithelial–stromal communication in the p38α uKO milieu.

Among ncWNTs, WNT5A plays a critical role in early pregnancy^27, 28^. WNT5A activates receptor tyrosine kinases ROR1 and ROR2 in the uterus. Both overexpression and deletion of WNT5A compromises pregnancy outcomes owing to sustained apicobasal polarity in the luminal epithelium^27^. Similar to WNT5A, IGF1 is also involved in epithelial depolarity and is highly expressed in the day 4 uterine stroma, activating IGF1R and downstream STAT3 in the luminal epithelia^38^. Uterine-specific deletion of IGF1R results in flawed embryo attachment because of sustained epithelial polarity^38^. These findings prompted us to examine these two molecules in the p38α uKO milieu. We first compared *Wnt5a* and *Igf1* expression using scRNA-seq data and found significantly upregulated *Wnt5a* and downregulated *Igf1* in both the luminal epithelium (LE) and decidualizing stroma (Dec) (Fig. 9a and b). As p38α has been implicated in the regulation of epithelial polarity, cytoskeletal organization, and E-cadherin localization, we next examined whether epithelial adhesion-and polarity-associated features were altered in p38α uKO uteri.

**Fig. 9.**
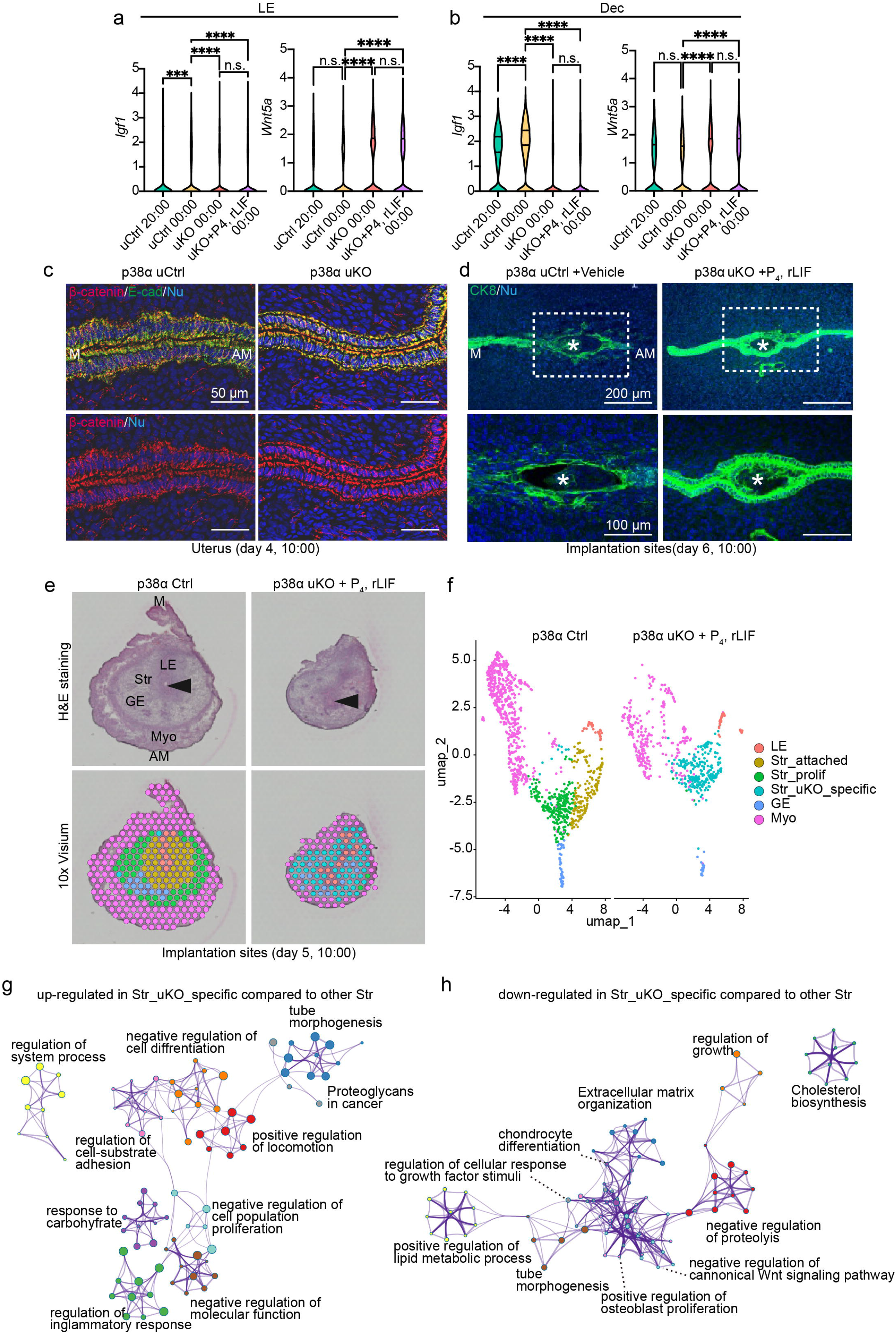
p38α uKO mice supplemented with P_4_ and LIF show embryo invasion failure associated with persistent epithelial polarity. **a, b** Luminal epithelial (**a**) and stromal (**b**) expression of *Igf1* and *Wnt5a* determined by scRNA-seq in each genotype and condition. \*\*\**P* < 0.001, \*\*\*\**P* < 0.0001, ns: not significant by one-way ANOVA followed by Bonferroni’s post-hoc test. **c** Representative images of immunostaining for β-catenin (red) and E-cadherin (green) in day 4 uteri from each genotype. M: mesometrial pole, AM: anti-mesometrial pole. Scale bar = 50 µm. **d** Representative images of immunostaining for CK-8 (green) at day 6 implantation sites from control and p38α uKO mice treated with P_4_ and rLIF. Areas demarcated by dashed lines in the top panels are shown in the bottom panels. M: mesometrial pole, AM: anti-mesometrial pole. Asterisks indicate embryos. Scale bar = 200 µm (top) and 100 µm (bottom). **e** H&E staining (upper) and visualization of the spatial transcriptome (lower) at day 5 implantation sites from the control and p38α uKO mice treated with P_4_ and rLIF. M: mesometrial pole; AM: anti-mesometrial pole, LE: luminal epithelia, GE: glandular epithelia, Str: stroma. Arrowheads indicate embryos. **f** UMAP analysis of the spatial transcriptome dataset colored according to cell type for day 5 implantation sites in control (left) and p38α uKO uteri treated with P_4_ and rLIF (right). Each dot color is colored according to the spatial transcriptome visualized in the uterine sections shown in (**e**). **g, h** Network of GO terms related to the upregulated (**g**) or downregulated genes (**h**) in Str_uKO_specific compared with the other stromal clusters. Each node represents an enriched term and is colored according to its cluster ID.

Immunostaining revealed stronger and more persistent localization of E-cadherin and β-catenin in the luminal epithelium of day 4 p38α uKO uteri compared with controls (Fig. 9c). These findings suggest that p38α loss is associated with abnormal maintenance or dysregulation of epithelial junctional-and polarity-associated states during the peri-implantation period. We then asked whether embryo invasion was influenced by p38α deletion. Our group has previously shown that epithelial removal after the loss of epithelial polarity facilitates embryo invasion and the subsequent pregnancy processes^15, 39^. To assess embryo invasion, the implantation sites were stained for cytokeratin 8 (CK8), a marker of epithelial and trophoblastic cells. In p38α uCtrl, the endometrial luminal epithelium disappeared and trophoblasts invaded the uterine stroma on the morning of day 6, whereas the endometrial luminal epithelium around the embryo remained and embryo invasion was incomplete in p38α uKO even after treatment with P_4_ and rLIF (Fig. 9d).

We also examined the spatial transcriptome of the embryo attachment site on the morning of day 5 from Ctrl-and P_4_ + rLIF-treated uKO mice to observe epithelial–stromal interactions (Fig. 9e–h). Cross-sectional tissues of each implantation site were clustered into six types: LE, embryo attached stroma (Str_attached), proliferative stroma (Str_prolif), uKO-specific Str (Str_uKO_specific), GE, and myometria (Myo) (Fig. 9e, f, Table S7). Notably, uKO tissues solely contained Str_uKO_specific as the stromal cluster. To understand the characteristics of this cell type, pathway analyses were performed using Metascape. We found that the upregulated genes in Str_uKO_specific were enriched in pathways such as “regulation of inflammatory response,” “negative regulation of cell differentiation,” and “negative regulation of cell population proliferation” (Fig. 9g, Table S8)

Down-regulated genes were related to “extracellular matrix organization,” “negative regulation of canonical Wnt signaling pathway,”, and “regulation of cellular response to growth factor stimuli” (Fig. 9h, Table S8), which are known to be associated with healthy pregnancy outcomes^37, 40^. These results indicate that embryo attachment to the uKO luminal epithelium induces inflammatory signals rather than physiological reactions in the stroma, compromising subsequent pregnancy maintenance. In summary, our data support an important role for uterine p38α in luminal remodeling, epithelial–stromal communication, and the establishment of appropriate embryo attachment.

## DISCUSSION

Communication between embryos and the endometrium is crucial for the successful establishment of pregnancy^41^. In preparation for blastocyst attachment, the endometrium undergoes massive morphological changes, especially in the epithelial layers: (1) PDS before embryo attachment, (2) slit formation of the endometrial lumen before embryo attachment, and (3) crypt formation of the endometrial luminal epithelium after embryo attachment. In this study, we conducted 3D histological analysis as well as single-cell and spatial transcriptome analyses to elucidate the details of dynamic morphological changes in the endometrial lumen. Two molecular pathways, the P_4_-PGR and LIF-STAT3 pathways, have so far been considered important for embryo attachment. Furthermore, we demonstrated a previously unappreciated role of p38α in addition to these two major pathways (Fig. 10).

**Fig. 10.**
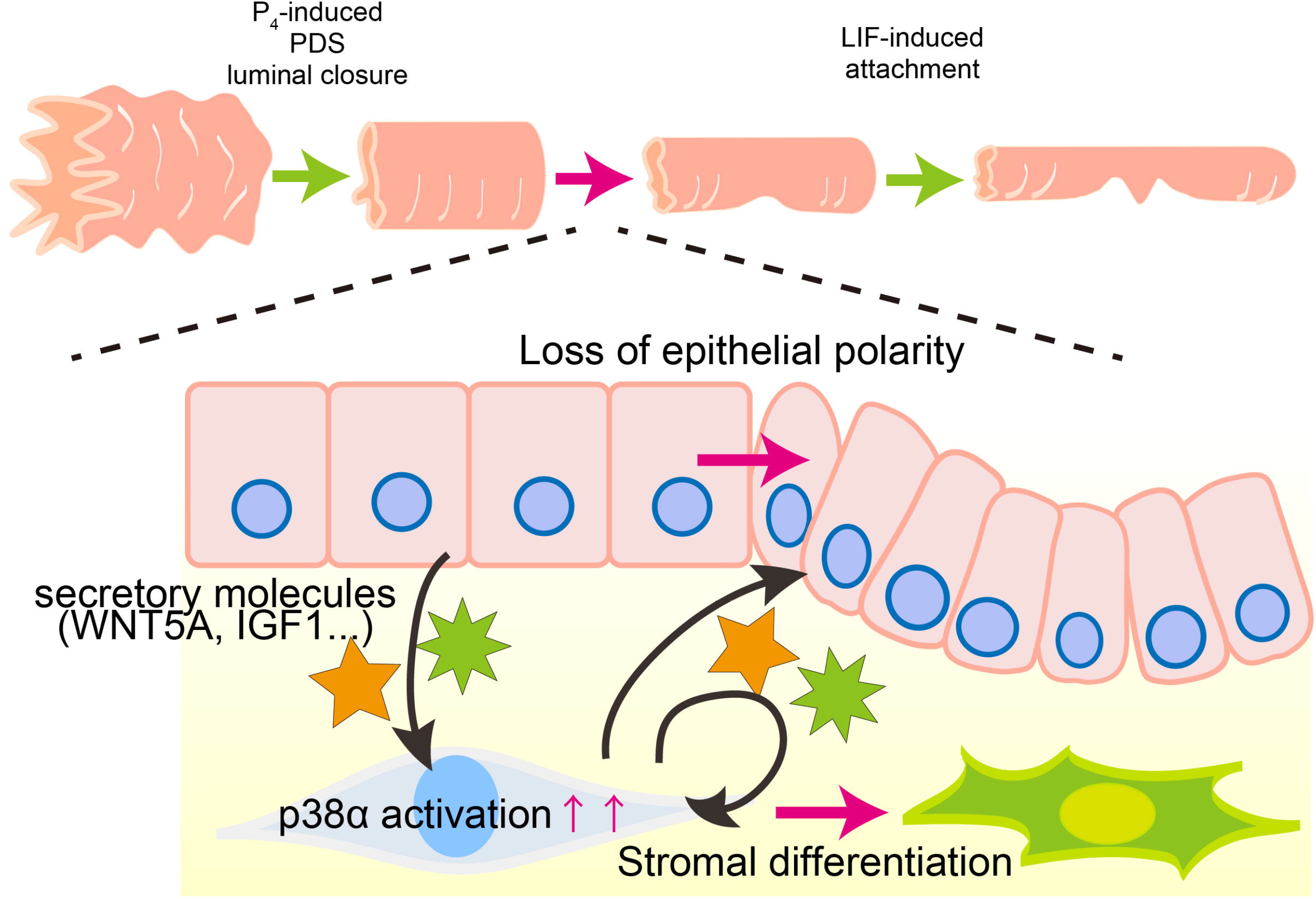
The role of uterine p38α in embryo attachment. p38α contributes to flattening of the endometrial luminal surface and induction of luminal narrowing prior to embryo attachment through pathways activated by P_4_. It is also suggested that uterine p38α induces glandular LIF, thereby promoting embryo attachment. In addition to these pathways, uterine p38α-dependent responses are associated with appropriate epithelial–stromal communication and remodeling of epithelial polarity-associated features, thereby supporting embryo attachment and subsequent invasion.

The major finding of this study was that luminal shapes were altered throughout the uterine horns during early pregnancy. In particular, changes were evident from the morning of day 4 to midnight. These morphological changes coincided temporally with embryo spacing and attachment. A previous study demonstrated that embryo spacing occurs from noon to evening on day 4, which was compromised in mice with systemic knockout of LPAR3, a lipid receptor expressed in the luminal epithelium^42^. Indeed, we observed that epithelial folding in the M–AM axis became evenly distributed, corresponding to the positions of the embryos from morning to evening on day 4, revealing a spatiotemporal association between luminal remodeling and embryo spacing. Previous studies have also implicated myometrial activity in embryo spacing^43^. Furthermore, a key strength of this study is the quantitative assessment of the relationship between epithelial folding and embryo positioning. By measuring the angle of luminal folds relative to the M–AM axis, we were able to evaluate the spatial relationship between folded and stretched luminal regions and the location of embryos. This relationship was clearly observed in control mice but was not detected in p38uKO mice. These findings suggest that, in p38uKO mice, persistent epithelial branching and failure of luminal flattening were associated with less stereotyped embryo positioning, making it difficult to accurately distinguish stretched and shrunken luminal regions or to evaluate the orientation of luminal folds.

This 3D imaging experiment had some limitations. As the data were obtained from euthanized mice, following changes over time in the same individual was impossible, and the influence of phenotypic time differences among individuals could not be completely ruled out. To examine epithelial morphology and phenotype over time in the same individual, an experimental system using techniques such as *in vivo* live imaging needs to be established.

An important point in interpreting the present genetic data is that *Pgr-*Cre deletes p38α in multiple uterine compartments, including epithelial and stromal cells. Therefore, the severe infertility phenotype in p38α uKO mice cannot be attributed solely to an epithelial-autonomous function of p38α. In this regard, our epithelial-specific *Ltf-*iCre deletion analysis is informative: epithelial deletion of p38α did not recapitulate the complete infertility or severe luminal disorganization observed in *Pgr*-Cre-mediated uKO mice. Together with our single-cell and spatial transcriptomic analyses showing abnormal stromal differentiation and disrupted epithelial–stromal signaling in p38α uKO uteri, these results support a model in which non-epithelial p38α-dependent responses and epithelial–stromal communication contribute to the establishment of implantation-competent luminal architecture.

In the present study, P_4_ and rLIF supplementation restored embryo attachment in p38α uKO mice; however, embryo invasion remained completely absent, resulting in subsequent pregnancy failure. Previous studies from our group have shown that uterine deficiency of HIF2α, Rb1, Ezh2, or Lox impairs embryo invasion and leads to pregnancy failure, highlighting the importance of coordinated epithelial–stromal remodeling during implantation^15, 16, 39, 44^. In the p38α uKO uteri, although P_4_ and rLIF supplementation restored key molecular signals required for attachment, abnormalities in luminal morphology persisted. Therefore, persistent defects in luminal remodeling may contribute to the failure of embryo invasion and subsequent pregnancy progression. However, because embryo attachment is severely impaired in untreated p38α uKO mice, the post-attachment role of stromal p38α cannot be directly assessed in the current model. Further studies using decidual cell-specific genetic approaches will be required to clarify the contribution of stromal p38α to embryo invasion.

Considering the functional features of p38α as a MAP kinase, the extracellular or intracellular stimuli that evoke p38α-dependent epithelial shaping remain unclear. MAP kinases can be activated downstream of various receptors, including receptor tyrosine kinases and G protein-coupled receptors^18^. One candidate is the ROR1/ROR2 receptor tyrosine kinase, which can be activated by WNT5A^27^. Uterine-specific deletion of WNT5A-ROR1/ROR2 axis resulted in defective embryo implantation because of abnormal luminal morphology, similar to our observations in p38α uKO mice. Similarly, deletion of IGF1-IGFR signaling causes poor embryo attachment, accompanied by abnormal luminal integrity^38^. These findings may explain why WNT5A and IGF1 were upregulated in p38α uKO uteri. Previous studies have shown that p38α regulates epithelial polarity, cytoskeletal organization, and E-cadherin localization in epithelial tissues. Consistent with this concept, we observed enhanced or persistent E-cadherin and β-catenin staining in the luminal epithelium of p38α uKO uteri. These results suggest that the abnormal luminal folding in p38α uKO mice may be linked to impaired remodeling of epithelial adhesion-and polarity-associated structures. Rather than indicating a simple loss of epithelial junctions, our findings suggest that p38α deficiency causes inappropriate persistence of epithelial junctional and polarity-associated features at a stage when the luminal epithelium should undergo dynamic remodeling to support embryo attachment and invasion. Importantly, as already mentioned, epithelial-specific deletion of p38α did not reproduce the severe phenotype observed in *Pgr*-Cre-mediated uKO mice. Thus, the altered epithelial adhesion/polarity phenotype in p38α uKO uteri is likely not solely due to epithelial-autonomous p38α loss but may also reflect abnormal stromal differentiation and disrupted epithelial–stromal communication.

In addition to altered epithelial adhesion and polarity, dysregulated epithelial transport and uterine fluid homeostasis may contribute to the abnormal luminal remodeling observed in p38α uKO uteri. Uterine fluid absorption is closely associated with luminal closure and remodeling during implantation^45, 46^. Our transcriptomic dataset revealed altered expression of several transport-and barrier-associated genes in p38α uKO uteri without P_4_ and rLIF supplementation, including *Aqp3*, *Slc26a6*, *Cldn10*, *Gjb2*, *Trpv6*, *Trpm6*, and *Atp6v0a4*. These genes are involved in water and ion transport, paracellular permeability, intercellular communication, and pH homeostasis, suggesting that dysregulated epithelial transport may contribute, at least in part, to the impaired luminal closure and remodeling in the mutant uterus. However, transcriptomic alterations alone do not demonstrate changes in transporter activity or uterine fluid dynamics. Direct measurements of uterine fluid volume and composition, together with functional analyses of epithelial transport, will therefore be required to determine whether dysregulated fluid absorption is a causal component of the implantation defects in p38α uKO mice.

New insights into the morphological changes in the endometrial lumen may provide an innovative approach to treating embryo attachment failure by focusing on endometrial lumen morphology. In humans, healthy implantation occurs at the fundus of the uterus^47^, which differs from that in rodents with tubular uterine structures. However, epithelial integrity is a common feature that regulates appropriate embryo implantation across species; for instance, after ovulation in humans, epithelial–mesenchymal transition with reduced epithelial polarity occurs during menstruation and implantation, thus influencing implantation outcomes^48, 49, 50^. A previous study using human endometrial epithelial cell lines demonstrated that deletion of p38α influenced the cellular transcriptome and metabolome, contributing to cancer cell-like characters^51^, suggesting a role for p38α in maintaining luminal epithelial integrity in humans. In addition to specific molecular mechanisms, implantation failure may also be overcome from the mechanical aspect of tissue mobility in pathological conditions such as uterine myoma and uterine adenomyosis, wherein abnormal morphological changes in the endometrial lumen are presumed^52^. These findings are expected to have broad applications in diagnosing and treating human implantation failure, including the search for biomarkers of implantation ability and supplementation with relevant molecules. As reported previously, p38α also plays critical roles in mammary gland lumen formation^22^; therefore, the mechanism discovered in this study could be applicable to other epithelial systems as well.

## METHODS

### Mice

WT (C57BL/6N, SLC), *p38*α-floxed (*Mapk14*-floxed; kindly provided by Dr. Kinya Otsu, University of Osaka)^19^, *Pgr-Cre*^53^, were used in this study. *Pgr-Cre* is expressed throughout the uterine layers^53^, and *Ltf*-iCre is expressed throughout the uterine epithelium ^54^. Mice with p38α deletion in all uterine layers (*Mapk14^flox/flox^ Pgr^Cre/+^*; p38α uKO) and deletion in the uterine epithelium (*Mapk14^flox/flox^ Ltf^Cre/+^*; p38α eKO) were generated by crossing either *Pgr-*Cre *or Ltf-*iCre with *p38*α-floxed mice, respectively. Cre-negative littermates (*Mapk14^flox/flox^*) served as controls. All mice used in this study were housed at the University of Tokyo Animal Care Facility, following the institutional guidelines for the use of laboratory animals.

### Evaluation of pregnancy outcomes

To examine the pregnancy outcomes, p38α-uKO, or p38α-floxed (control) female mice were mated with fertile C57BL/6N male mice, as reported in previous studies^15, 39, 55^. The day of vaginal plug detection was considered day 1 of pregnancy. Pregnant mice were euthanized by cervical dislocation on the designated day of pregnancy to evaluate pregnancy phenotypes and collect samples. On days 2 and 3, both oviducts were flushed with saline to confirm the presence of 2-cell embryos on day 2 and 8-cell embryos or morulae on day 3. On day 4, one uterine horn was flushed with saline to confirm the presence of blastocysts. Embryo attachment sites were observed as blue bands soon after intravenous injection of a 1% solution of Chicago blue dye (Sigma-Aldrich) in saline on days 5 and 6. When no embryo attachment sites were observed on day 5, both uterine horns were cut and flushed with saline to collect the embryos.

To analyze implantation failure, pregnant mice were sacrificed on day 8 at 10:00, and implantation sites were histologically assessed. If obvious hematopoietic cell infiltration was observed, implantation failure was diagnosed. Parturition events were monitored daily from days 19 to 22. All mice were then dissected, and the abdominal cavity was observed for evidence of miscarriage.

To evaluate the phenotype of embryo attachment in wild-type mice over time, we observed the uteri on day 4 morning (day 4 10:00), day 4 evening (day 4 16:00), day 4 night (day 4 20:00), day 4 midnight (day 5 00:00), and day 5 morning (day 5 10:00). For day 5 night, the observation time was defined as 18:00–22:00. In preliminary experiments, we confirmed that the phenotypes were equivalent, with no significant changes in the morphology of tissues collected during this 4-h period. As described above, samples with visible embryo attachment sites were evaluated for the number of attachment sites, whereas those with no visible embryo attachment sites were evaluated as pregnant specimens by flushing the contralateral uterine horn with saline and observing the blastocysts.

Daily subcutaneous injections of P_4_ (2 mg/mouse/day) were administered to p38α uKO mice from day 2 of pregnancy or from the designated day of pregnancy at 10:00, as previously described^15^.

rLIF injections were administered as previously reported^11^. Female mice received rLIF (20 µg/head, i.p.) at 9:00 and 18:00 on day 4 of pregnancy. The rLIF expression vector was a kind gift from Prof. Eichi Hondo^13^.

### RNA extraction and real time-quantitative PCR (RT-qPCR)

RNA was prepared from homogenized frozen tissues, as previously described^16^. qRT-PCR was performed using THUNDERBIRD SYBR qPCR Mix (TOYOBO). The housekeeping gene *Actb* was used for internal standardization of mRNA expression. Relative expression levels were determined using the ΔΔCt method^56^. The following primers were used.

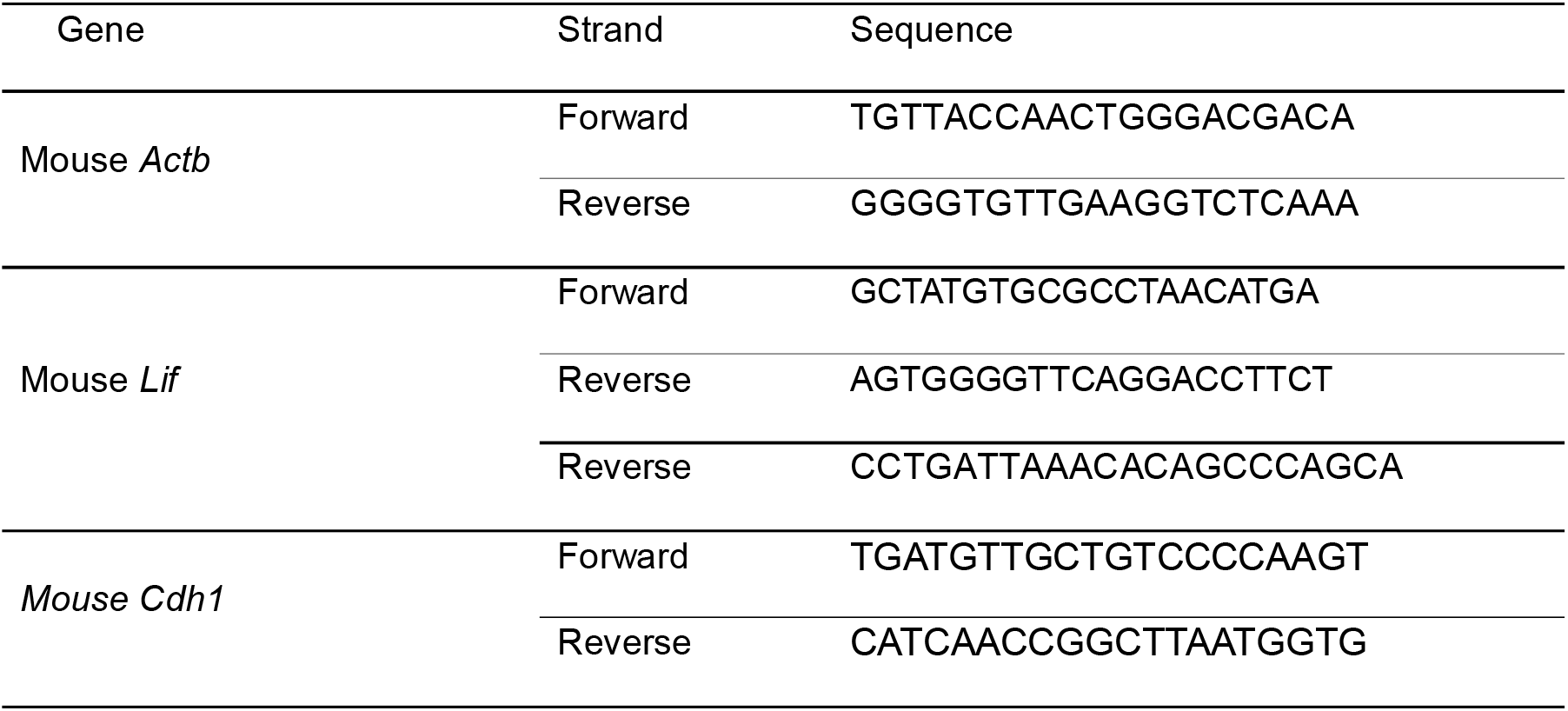

### H&E staining and immunostaining

H&E staining and immunostaining of uterine tissues were performed using paraffin-embedded sections (6 μm) or frozen sections (12 μm), as previously described^15^. For immunohistochemistry, the sections were incubated overnight with primary antibodies, including p38 (8690, Cell Signaling Technology, 1:800), pp38 (4511, Cell Signaling Technology, 1:800), ESR1 (ab32063, Abcam, 1:200), PGR (ab63605, Abcam, 1:100), pSTAT3 (ab76315, Abcam, 1:100), FOXA2 (8186, Cell Signaling Technology, 1:200), and COX2 (AA570-598, Cayman,1:200). Signals were detected using a DAB substrate kit (#425011, Nichirei) after incubation with horseradish peroxidase-conjugated secondary antibodies (K4003, Dako). Images were captured using a Leica DM5000 B light microscope.

For immunofluorescence analysis of paraffin sections, the sections were incubated overnight with primary antibodies, including CK8 (DSHB, 1:500), and signals were detected using Alexa Fluor 488-conjugated anti-rat immunoglobulin G (Thermo Fisher Scientific, A11006,1:500). Nuclei were stained with 6-diamidino-2-phenylindole (DAPI) (Dojindo, 1:500). For immunofluorescence analysis of frozen sections, the sections were incubated overnight with primary antibodies, including Ki67 (20701, Cell Signaling Technology, 1:200, Alexa Fluor® 555 Conjugate), E-cadherin (3199, Cell Signaling Technology,1:200, Alexa Fluor® 488 Conjugate), and β-catenin (83539, Cell Signaling Technology, 1:200, Alexa Fluor® 555 Conjugate). Nuclei were detected using 6-diamidino-2-phenylindole DAPI (1:500). Images were captured using an AXR microscope (Nikon).

### Automated western blots using Simple Western (WES)

Proteins were extracted from cryopreserved and homogenized day 4 uterine tissues using RIPA buffer (Sigma) supplemented with a proteinase inhibitor cocktail (Sigma) and a phosphatase inhibitor cocktail (Sigma).

Equal amounts of protein (2 µg/µL) were loaded into a 12–230 kDa Separation Module Kit and analyzed using the Protein Simple Wes® System (Protein Simple, San Jose, CA, USA), following the manufacturer’s instructions. The antibodies used included Actin (C-11, sc-1615, Santa Cruz, 1:2000), STAT3 (4904, Cell Signaling Technology, 1:2000), and pSTAT3 (9145, Cell Signaling Technology, 1:2000). Anti-goat IgG and anti-rabbit IgG antibodies were used as secondary antibodies. Actin served as the loading control.

### 3D visualization of uterine endometrial luminal epithelium

3D visualization of the day 1–4 uteri or day 5 and 6 implantation sites was performed, as previously reported^10^. Samples were fixed in Dent’s fixative (methanol: DMSO = 4:1) overnight at –20 °C and then washed with 100% methanol. The samples were bleached with 3% H_2_O_2_ in methanol at 4 °C overnight to remove pigmentation. After washing in PBST six times for 1 h each, the samples were incubated with an anti-E-cadherin antibody (Cell Signaling Technology, 24E10, 1:500) at room temperature on a rotator for 5–7 days. Following incubation, the samples were washed in PBST six times for 1 h each at room temperature. The samples were then incubated with an Alexa Fluor 555-conjugated anti-rabbit secondary antibody (A21428, Thermo Fisher Scientific; 1:500 dilution) at room temperature on a rotator for 5–7 days in the dark. After washing in PBST six times for 1 h each at room temperature, the samples were fixed in methanol for 30 min at room temperature. Finally, the tissues were cleared in BABB (benzyl alcohol: benzyl benzoate = 1:2) at room temperature for more than 1 h in the dark. Z-stack confocal 3D images were acquired using an AXR microscope (version 5.30; Nikon). Images were acquired using a 10× objective lens with a 3-µm Z-stack interval. To visualize the adenomyosis lesions in 3D, the Surface Tool in IMARIS (version 9.8, Oxford Instruments) was used. Surface reconstruction of the E-cadherin regions was performed using automatic thresholding, enabling visualization and quantitative analysis of tissue morphology.

### Quantitative analysis of luminal epithelial changes

As previously reported, ^31^, to quantify the transition between shrunken and stretched areas in wild-type mice before and after embryo attachment, the angle of luminal folding against the M–AM axis was quantified at the day 4 20:00 and day 5 00:00 time points. For each implantation site, the angle between the gland directly associated with the embryo and the longitudinal axis of the uterus was measured using ImageJ at three locations: the center of the embryo and positions ±30 μm from the embryo center. The mean value of the three measurements was used for quantitative analysis.

To quantitatively evaluate epithelial flattening morphology, three representative 2D cross-sectional images were generated from each 3D reconstructed dataset along the longitudinal (x) axis. Sections were collected at the midpoint of the image stack and at positions ±500 μm from the midpoint. Epithelial staining area and epithelial branching features were subsequently quantified from these cross-sectional images using identical analysis parameters across samples.

### Measurement of serum E₂ and P_4_ levels

Blood samples were collected from mice on the indicated day of pregnancy. Serum P_4_ levels were measured, as described previously^39^, using a progesterone enzyme-linked immunosorbent assay (ELISA) kit (582601, Cayman). Serum E₂ levels were measured using an estradiol ELISA kit (501890, Cayman).

### Spatial transcriptomics

Spatial transcriptomes were analyzed using the 10x Visium (10x Genomics) following the manufacturer’s protocol. Day 5 morning (10:00) uteri from control and p38α uKO females were collected. Frozen sections (10 μm) were mounted on gene expression slides and sent to KOTAI Bio Inc. (Osaka, Japan) for processing. Following a 30-min Proteinase K treatment, the sections were hybridized with spatial tags on the slides and reverse-transcribed *in situ*. The cDNAs were analyzed by RNA sequencing using DNBseq (MGI) with 300 million reads per sample. Raw FASTQ files and microscope slide images for each sample were processed with Space Ranger software (version 1.1, 10× Genomics) using the “spaceranger count” pipeline, involving STAR with the default parameters for aligning reads against the mouse reference genome mm10 “refdata-gex-mm10-2020-A.” This pipeline uses Visium spatial barcodes to generate a feature spot matrix with unique molecular identifier counts. Clustering analysis was performed using Seurat (version 5.0.0)^57^, and clusters were visualized using UMAP. Differentially expressed genes between genotypes were identified using an adjusted *P* value < 0.05 and a fold change > 1.5. Metascape^34^ and Enrichr^58^ were used to analyze the GO terms and upstream transcription factors within each cluster, respectively. The mouse Visium data were deposited in the GEO database (Accession No. GSE305995).

### scRNA-seq and data analysis

The 10x Genomics Chromium FRP protocol was followed for scRNA-seq analysis. On days 4 and 5 for WT mice, or on day 4 evening and midnight for p38α-floxed and p38α uKO mice, uterine horns were excised, snap-frozen, and sent to Takara Bio Co. (Osaka, Japan). After cell fixation with formaldehyde, a single-cell suspension was prepared using a gentleMACS (Miletenyi Biotec). The cells were used for RNA sequencing library preparation using the Chromium Next GEM Single Cell Fixed RNA Sample Preparation Kit, Chromium Fixed RNA Kit, Mouse Transcriptome, Chromium Mouse Transcriptome Probe Set v1.0.1, Chromium Next GEM Chip Q Single Cell Kit, Dual Index Kit TS Set A, and Chromium X (10x genomics, USA). Paired-end sequencing was performed on an Illumina next-generation sequencer (NovaSeq 6000; Illumina). Raw FASTQ files were processed using Cell Ranger software (10x Genomics, USA), and Seurat (http://www.satijalab.org/seurat) v5.0.0^57^ was used to process read counts. Cell trajectories were determined using Monocle3^59^. CellChat was used for the cell–cell interaction analysis. The mouse scRNA-seq data were deposited in the GEO database (Accession Nos. GSE296583 and GSE305994).

## Statistical analyses

Statistical analyses were performed using a two-tailed Student’s *t*-test or one-way analysis of variance (ANOVA), followed by Bonferroni post-hoc tests, in GraphPad Prism10. Statistical significance was set at *P* < 0.05.

## Study approval

All animal experiments were approved by the Institutional Animal Experiment Committee of the University of Tokyo Graduate School of Medicine (approval numbers P20-076 and A2023M165).

## Data and material availability

The RNA-seq experimental data will be made publicly available upon publication (GSE305994 and GSE305995). This study did not involve the development of custom code or algorithms. All the software used in this study is publicly available and is cited in the main text and the Methods section.

## Acknowledgments

We thank Ms. Atsumi Miura for providing technical assistance. We are grateful to Francesco J. DeMayo (National Institute of Environmental Health Sciences) for providing *Pgr*-Cre mice, and to Kinya Otsuki (Osaka University) for providing *Mapk14*-floxed mice.

## Funding

This work was supported by the Japan Society for the Promotion of Science (JSPS) KAKENHI (grant nos. 23K08278, 23K27176, 24K22157, 24K21911, 25K02779, and 25H01065), Japan Agency for Medical Research and Development (AMED) (grant nos. JP25gn0110085, JP24gn0110069, JP25gk0210039, JP24lk0310083, JP25gn0110097, JP25gk0210042, JP25gk0210045, and JP26gn01100111), Children and Families Agency (grant no. JPMH23DB0101), Japan Science and Technology Agency (JST) Fusion Oriented Research for Disruptive Science and Technology (FOREST) (grant no. JPMJFR210H), Mochida Memorial Foundation for Medical and Pharmaceutical Research, Uehara Memorial Foundation, Inoue Foundation for Science, Astellas Foundation for Research on Metabolic Disorders, The Naito Foundation and the fund for joint research with NIPRO Corporation.

## Author contributions

Conceptualization: S.A. and Y.H.; Funding acquisition: S.A. and Y.H.; Investigation: C.I., S.A., Y.F., X.H., R.S.H., D.H., T.H., M.M.; Data analysis: C.I., S.A.; Data interpretation: C.I., S.A., Y.H.; Project administration and supervision: Y.H.; Writing - original draft preparation: C.I., S.A.; Writing – review and editing: S.A.

## Competing interests

All authors declare that they have no competing interests.

## Notes

### Competing Interest Statement

The authors have declared no competing interest.

### Summary of Updates

This version of the manuscript has been revised to clarify the interpretation of uterine luminal remodeling during embryo implantation. The title, Abstract, Results, Discussion, and Figure 10 were updated to reflect new epithelial-specific knockout data showing that epithelial p38alpha loss alone is insufficient to reproduce the severe implantation phenotype observed in the uterine-wide knockout. The manuscript now places greater emphasis on uterine p38alpha-dependent responses and epithelial-stromal communication. Statements concerning embryo positioning and spacing were revised to avoid implying causal relationships that are not directly established by the time-course analyses. Quantitative analyses of luminal morphology were also clarified so that individual animals, rather than individual tissue sections, are defined as biological replicates. The interpretation of epithelial polarity and embryo invasion was further refined, including revision of the Figure 9 title and related text. Figure legends and Methods were updated for clarity and consistency, and labeling errors and terminology were corrected throughout the manuscript.

