## Supplementary Fig. for "A stress-responsive morphogenetic program coordinates uterine luminal remodeling during early pregnancy"

### Supplementary Figures

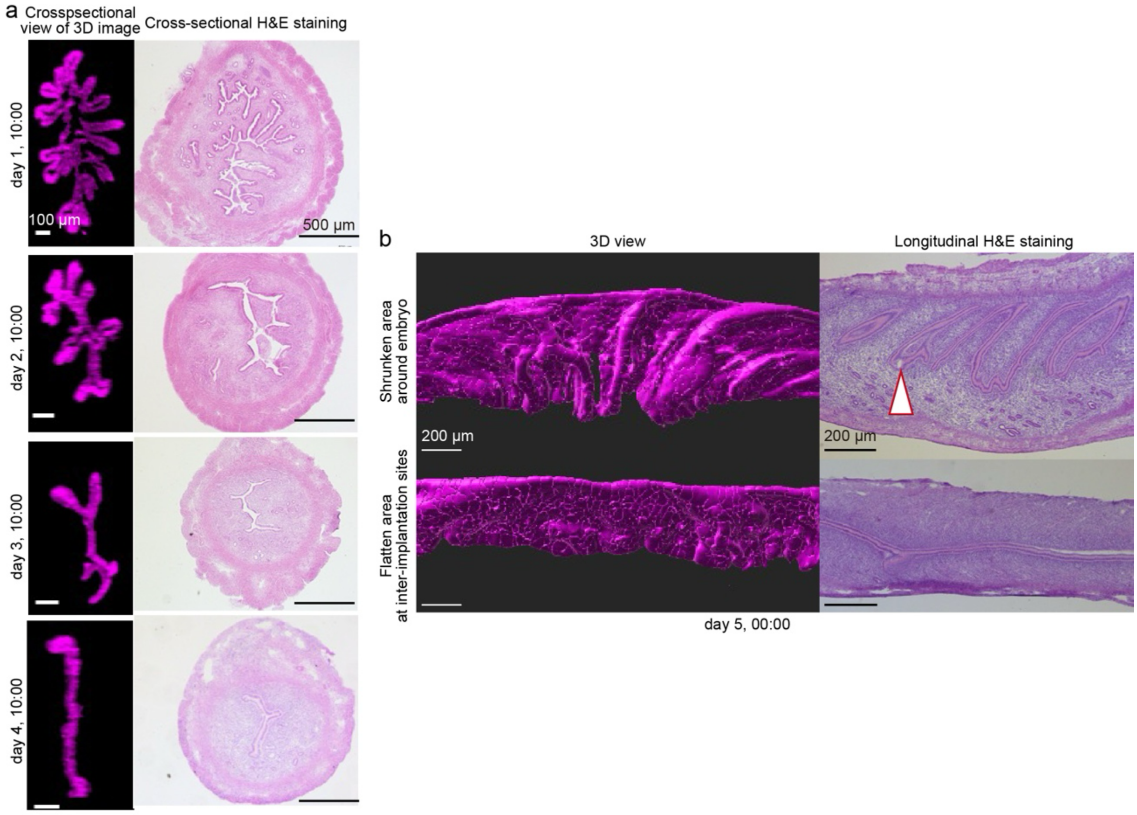


**Fig. S1 Validation of three-dimensional (3D) luminal imaging by comparison with H&E staining images.**

a. Representative images of cross-sectional views of 3D images (left) and H&E staining (right) for wild-type uteri on days 1–4 of pregnancy. Scale bar = 100 µm (left) and 500 µm (right).
b. Representative images of 3D views (left) and longitudinal H&E staining (right) for wild-type uteri on day 4 midnight. Scale bar = 200 µm.

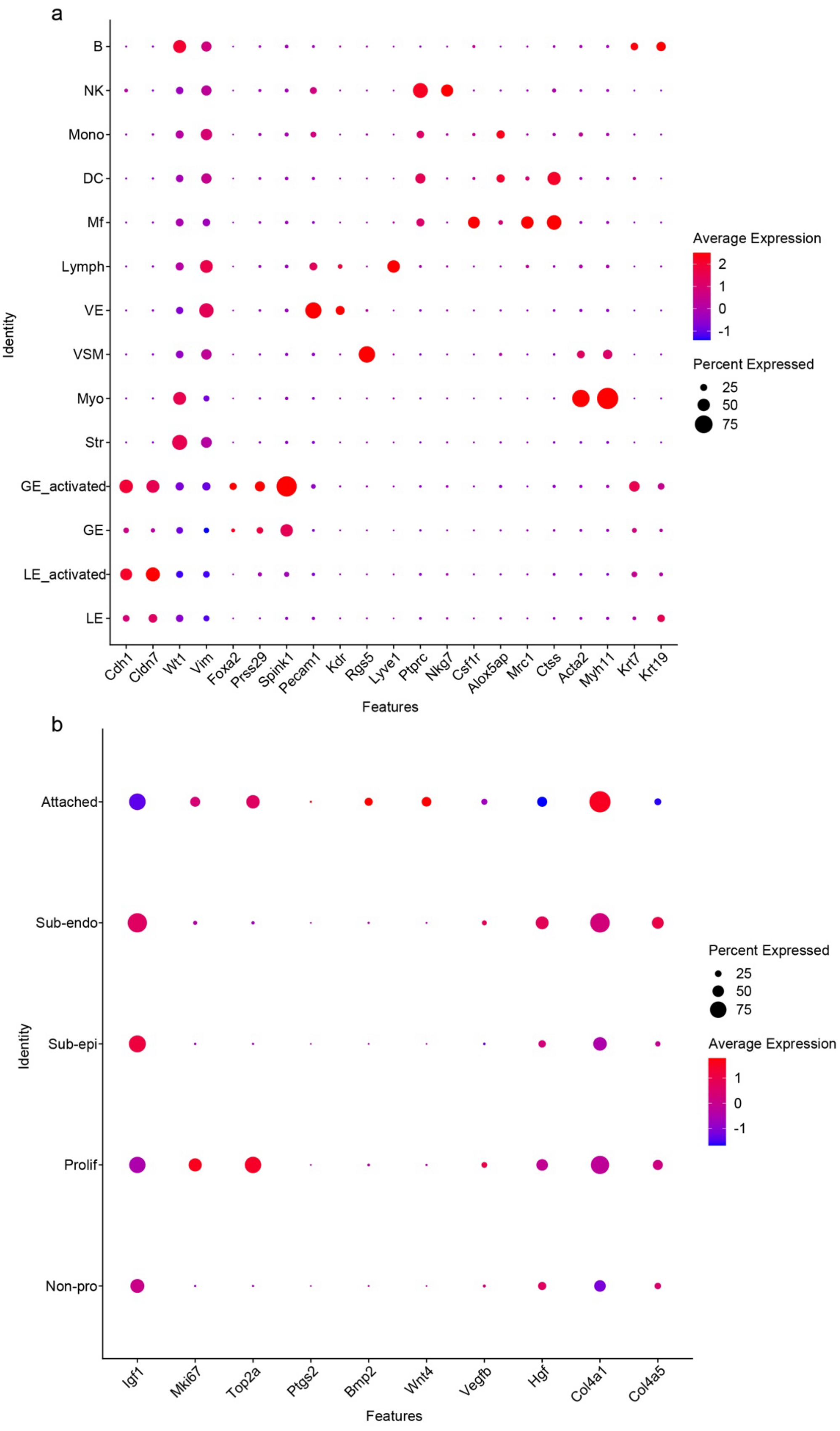

a
B
NK
Mono
DC
Mf
Lymph
VE
VSM
Myo
Str
GE_activated
GE
LE_activated
LE
Identity
Cdh1
Cldn7
Wt1
Vim
Foxa2
Prss29
Spink1
Pecam1
Kdr
Rgs5
Lyve1
Ptprc
Nkg7
Csf1r
Alox5ap
Mrc1
Ctss
Acta2
Myh11
Krt7
Krt19
Features
Average Expression
2
1
0
-1
Percent Expressed
25
50
75
b
Attached
Sub-endo
Sub-epi
Prolif
Non-pro
Identity
Igf1
Mki67
Top2a
Ptgs2
Bmp2
Wnt4
Vegfb
Hgf
Col4a1
Col4a5
Features
Percent Expressed
25
50
75
Average Expression
1
0
-1

**Fig. S2 Validation of each cell cluster using the corresponding marker gene expression.**

a. Dot plot showing the expression of marker genes in scRNA-seq datasets from mouse endometria on days 4–5. Dot color intensity represents the average expression from low (blue) to high (red), and dot size represents the percentage of cells in each cluster expressing that gene.

b. Dot plot showing the expression of stromal cell cluster marker genes in mouse endometria on days 4–5. Dot color intensity represents the average expression from low (blue) to high (red), and dot size represents the percentage of cells in each cluster expressing that gene.

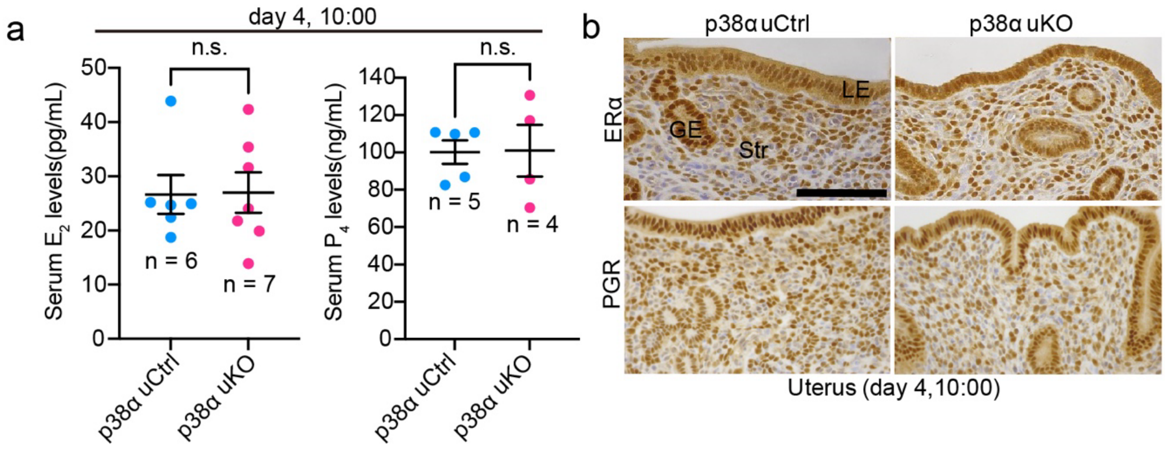


**Fig. S3 Comparable levels of female sex hormonal signaling between each genotype.**

a. Serum $E_2$ (left) and $P_4$ (right) levels from each genotype were tested on day 4 of pregnancy. The number of replicates is shown on each graph. Data represents the mean ± SEM, n.s.: not significant by Student's *t*-test.

b. Representative images of immunohistochemistry for ERα and PGR in day 4 uteri from each genotype. Scale bar = 100 μm, LE: luminal epithelia, GE: glandular epithelia, Str: stroma.

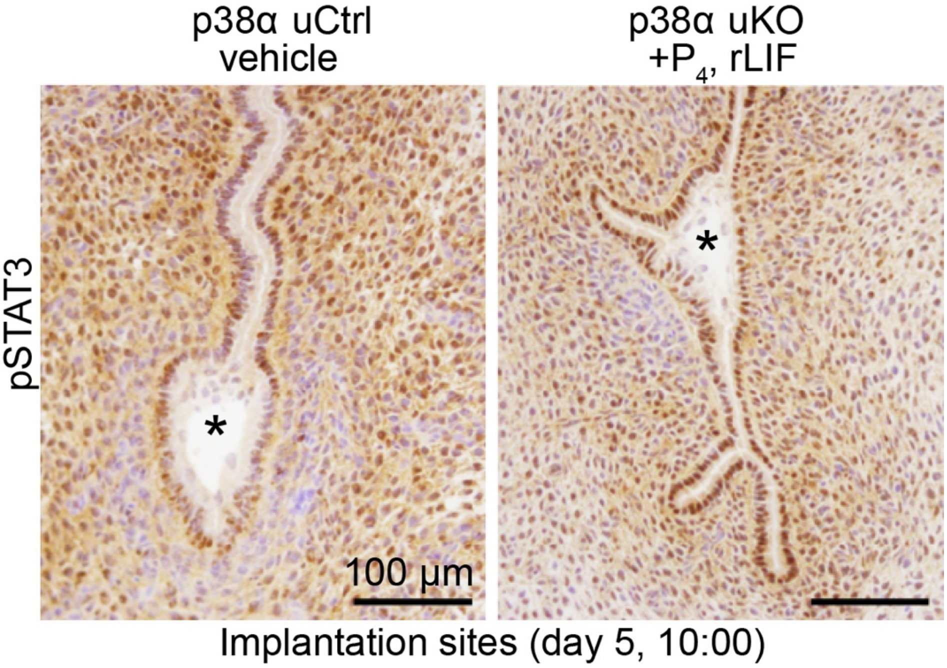


**Fig. S4 $P_4$ and rLIF supplementation in uKO induced embryo attachment, as depicted by pSTAT3 activation.**

Representative images of pStat3 on day 5 implantation sites from control or uKO mice treated with $P_4$ and rLIF. Scale bar = 100 μm. Asterisks indicate blastocysts.

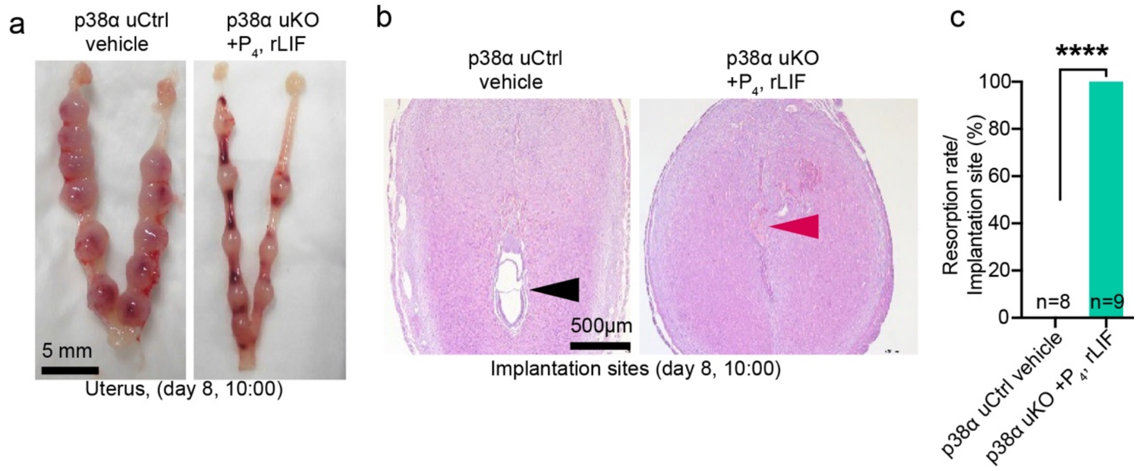


**Fig. S5 $P_4$ and rLIF supplementation could not restore pregnancy maintenance in uKO.**

a. Representative images of day 8 uteri from control or uKO mice treated with $P_4$ and rLIF. Scale bar = 5 mm.

b. Representative images of day 8 implantation sites from control or uKO mice treated with $P_4$ and rLIF. Scale bar = 500 μm. The black arrowhead indicates an embryo while the red arrowhead indicates a resorbed embryo.

c. The ratio of resorption in control or uKO mice treated with $P_4$ and rLIF on day 8 of pregnancy. The number of replicates is shown on each graph. ****$P < 0.0001$ by Fisher's test.

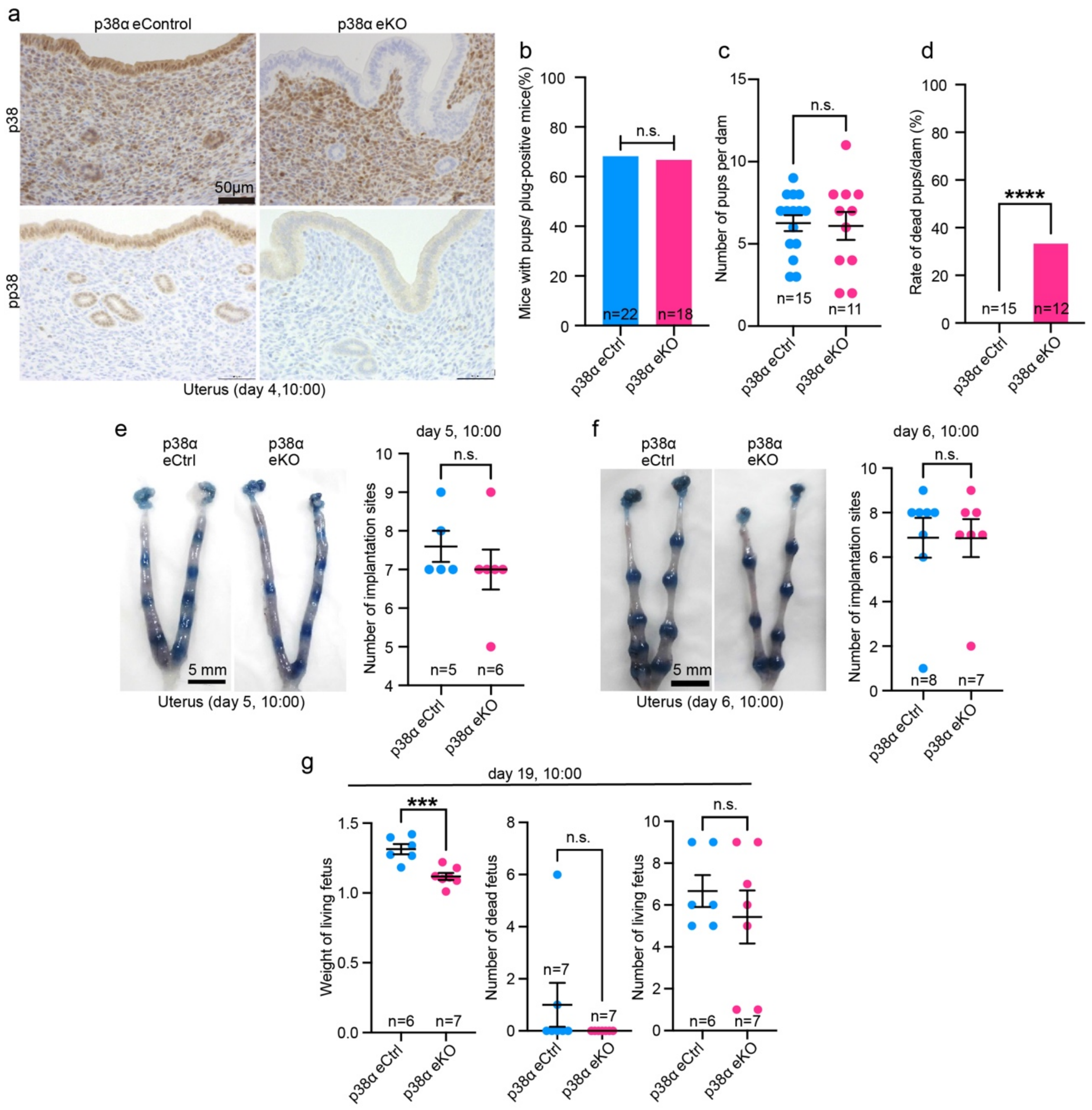


**Fig. S6 Comparable pregnancy outcomes in p38α eKO females compared with controls.**

a. Efficient deletion of p38α (top) and pp38α (bottom) specifically in the epithelium, as depicted by immunohistochemistry on day 4 of pregnancy. Scale bar = 50 μm.

b–d. Comparable levels of delivery rates (b) and litter sizes (c) in dams of each genotype, although the number of dead pups was increased in eKO (d). The number of replicates is shown on each graph. Data represents the mean ± SEM, n.s.: not significant, ****$P <$ 0.0001 by Student's t-test.

e, f. Representative images of uteri on day 5 (e, left) and day 6 (f, left). The average numbers of implantation sites are shown to the right of each image. Scale bar = 5 mm. The number of replicates is shown on each graph. Data represents the mean ± SEM, n.s.: not significant by Student's t-test.

g. The average weighs of living fetuses (left), numbers of dead fetuses (middle), and numbers of living fetuses (right) on day 19 of pregnancy were comparable between the genotypes. The number of replicates is shown on each graph. Data represent the mean ± SEM, n.s.: not significant, ***$P$ < 0.001 by Student's t-test.

**Supplementary Tables (Separate files)**

**Table S1 Marker genes of each cell cluster in whole endometrial tissues from mouse scRNA-seq data during days 4 and 5 of pregnancy.** See also Fig. 3a.

**Table S2. Upregulated genes in the LE_activated cluster compared with the LE cluster in mouse scRNA-seq data during days 4 and 5 of pregnancy.** See also Fig. 3b and c.

**Table S3 Marker genes of each cell cluster in stromal cells from mouse scRNA-seq data during days 4 and 5 of pregnancy.** See also Fig. 3d.

**Table S4. Upregulated genes in the Attached cluster compared with the other stromal clusters in mouse scRNA-seq data during days 4 and 5 of pregnancy.** See also Fig. 3e and f.

**Table S5. Marker genes of each cell cluster in whole endometrial tissues from scRNA-seq data comparing control and p38α uKO mice.** See also Fig. 8a.

**Table S6. Marker genes of each cell cluster in stromal cells from scRNA-seq data comparing control and p38α uKO mice**. See also Fig. 8b.

**Table S7. Marker genes of each cell cluster in whole endometrial tissues from mouse 10x Visium data comparing control and p38α uKO mice.** See also Fig. 9e.

**Table S8. DEGs in Str_uKO and other stromal clusters in 10x Visium data comparing control and p38α uKO mice.** See also Fig. 9g and h.
